# Abundant and active community members respond to the diel cycle in hot spring phototrophic mats

**DOI:** 10.1101/2024.08.19.608644

**Authors:** Amanda N. Shelton, Feiqiao B. Yu, Arthur R. Grossman, Devaki Bhaya

## Abstract

Photosynthetic microbial mats in hot springs can provide insights into the diel behaviors of communities in extreme environments. In this habitat, photosynthesis dominates during the day, leading to super-oxic conditions, with a rapid transition to fermentation and anoxia at night. Multiple samples were collected from two springs over several years to generate metagenomic and metatranscriptomic datasets. Metagenome assembled genomes (MAGs) comprised 71 taxa (in 19 different phyla), of which twelve core taxa were present at high abundance in both springs. The eight most active taxa identified by metatranscriptomics were an oxygenic cyanobacterium (*Synechococcus* sp.), five anoxygenic phototrophs from three different phyla, and two understudied heterotrophs from phylum Armatimonadota. Surprisingly, in all eight taxa, a significant fraction of genes exhibited a diel expression pattern although peak timing varied considerably. The two abundant heterotrophs exhibit starkly different peak timing of expression, which we propose is shaped by their metabolic and genomic potential to use carbon sources that become differentially available during the diel cycle. Network analysis revealed pathway expression patterns that had not previously been linked to diel cycles, including ribosome biogenesis and chaperones. This provides a framework for analyzing metabolically coupled communities and the dominant role of the diel cycle.

## Introduction

The microbial mats in the hot springs of Yellowstone National Park, USA, (YNP) have been referred to as the “tropical rainforests’’ of the microbial world because they harbor dense, diverse, stratified microbial communities dominated by phototrophs that use light energy to fix both carbon and nitrogen^1^. Octopus Spring (OS) and Mushroom Spring (MS) are two geographically close, circumneutral hot springs that have been extensively studied with a focus on the identification of phototrophic organisms and their abundance as a function of temperature ^2^. As water flows away from the spring source, a year-round stable temperature gradient is formed along the outflow channels and benthic microbial communities with varying compositions are found at different temperatures ^3^. Typically, stratified biofilms are formed between 50°C and 70°C, with the top green layer (1-2 mm) dominated by oxygenic cyanobacteria while the orange “undermat” layer harbors mostly anoxygenic phototrophs and less well-characterized heterotrophs ^4–6^. Light intensity, quality, and oxygen concentrations vary dramatically with mat depth, as does community composition ^4,7–12^. During the day, oxygenic photosynthesis carried out by the unicellular cyanobacterium, *Synechococcus* sp., (recently renamed *Thermostichus* sp.)^13^ creates a hyperoxic environment while carbon fixation causes an increase in the pH of the mat ^9,14^. However, as evening progresses, the rate of aerobic respiration exceeds the rate of photosynthetic O_2_ evolution, and the mats rapidly become anoxic. At night, cyanobacteria switch to fermentation of stored glycogen and with an accompanying decrease in pH^7–9^. Filamentous anoxygenic phototrophs such as *Roseiflexus* spp. and *Chloroflexus* spp. abundant in the undermat also fix carbon in the morning although they use reductants, such as organic acids or hydrogen, produced by other mat organisms ^15–17^.

Methods such as denaturing gradient gel electrophoresis (DGGE) and 16S ribosomal DNA amplicon sequencing were used to correlate “ecotypes” of abundant phototrophs with parameters such as temperature, light and oxygen, leading to the hypothesis that particular “ecotypes’’ have evolved to dominate in specific environmental niches ^3,12,18,19^. The advent of genomics, metagenomics and (meta)transcriptomics, coupled with physiological studies led to the identification of new taxa, and identified diel patterns of expression of sentinel genes of major metabolic pathways *in situ,* such as photosynthesis, fermentation, and nitrogen fixation. These studies also revealed complex patterns of genomic diversity and extensive recombination within the mat populations and provide a working framework or ‘snapshots of activity’ for major mat organisms. However, gaps remain in our understanding of community interactions, particularly those that are influenced by diel cycles ^9,18,20–25^.

We reasoned that coupling available biogeochemical data with extensive meta-omics analyses could provide insights into the diel behaviors of communities in these extreme environments ^3,7,8,18,21,22,26–28^. In this study, we used samples collected from OS and MS over several years and across a temperature gradient ranging from 50°C to 65°C for metagenomic analyses. We also analyzed metatranscriptomic data collected every two hours over a diel cycle and show that such an approach provides testable hypotheses and insights into some of the coordinated behavior of phototrophic communities.

We created a census of mat organisms and identified seventy-one taxa from nineteen phyla (species-to-genus level grouping), of which twelve were considered “core taxa” due to their high abundance in all samples while the rest were more variable in their abundance. MS and OS have somewhat different community structures while sharing the same abundant and active taxa. Eight taxa were identified as most active in the metatranscriptome datasets of which six were phototrophs. The remaining two active taxa were heterotrophs belonging to the understudied phylum Armatimonadota. Among the eight active taxa, a significant fraction of genes in each taxon exhibited a diel expression pattern but with several differences in the peak expression timing between taxa. Despite these significant diel differences, we were surprised to find that ribosome biogenesis appeared to peak in the morning in seven of the eight taxa.

Differences in expression timing of key pathways suggests that the two Armatimonadota taxa have evolved to use different carbon sources released by the phototrophs at different times in the diel cycle. These analyses help create a more comprehensive view of the phototropic mat community interactions and the dominant role of the diel cycle.

## Results and Discussion

### Extensive metagenomic sequencing recovers diverse taxa from Mushroom and Octopus Spring

To overcome the limited analytical power typically associated with access to a few samples and/or low sequence coverage, we carried out extensive metagenomic sequencing on the top green layer of mat cores collected from either OS or MS over 5 years (2004-2009), mostly from a 60°C site. Samples were taken from 4 different temperatures in both OS and MS in 2006 and 2009 (**Table 1**, **Fig. 1**, **Fig. 2, Supplementary Data 1**). A few samples were also taken from the thick orange undermat (top green layer removed manually) in 2004 and 2007 at 55°C and 60°C (labeled “um” in **Table 1**, **Fig. 2**, **Supplementary Data 1**).

**Figure 1:**
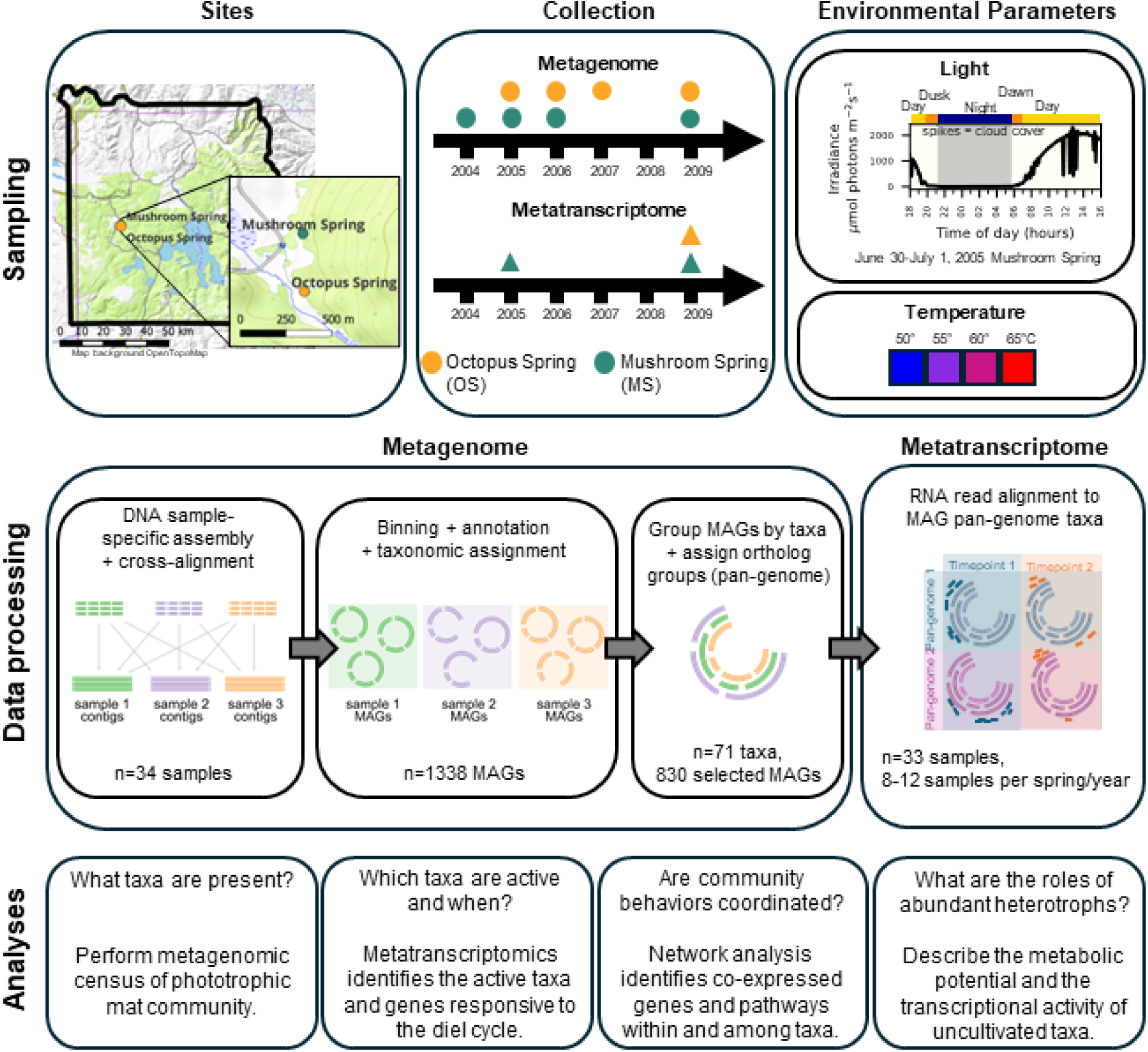
Overview of study site and workflow. Top panels: Sampling. Top left: Mat cores were collected from Mushroom and Octopus Springs in Yellowstone National Park during the years listed for metagenomics and metatranscriptomics (top middle). **Table 1** and **Supplementary Data 1** list more specific details. Top right: Environmental parameters for sampling. For light, a typical light curve for Mushroom Spring during our metatranscriptome sampling is depicted (adapted from Jensen *et al.* 2011). Middle panels: Data processing: Following DNA extraction, assembly, cross-alignment and binning, the anvi’o pan-genome pipeline was used to aggregate the selected MAGs into species-to-genus level taxa groups and to filter out MAGs of very-low quality or present in only a single sample. 830 MAGs were retained for further analyses, including for the RNA alignment for the metatranscriptome datasets. Bottom panels: Analyses performed to understand the mat community composition and activity.

**Figure 2:**
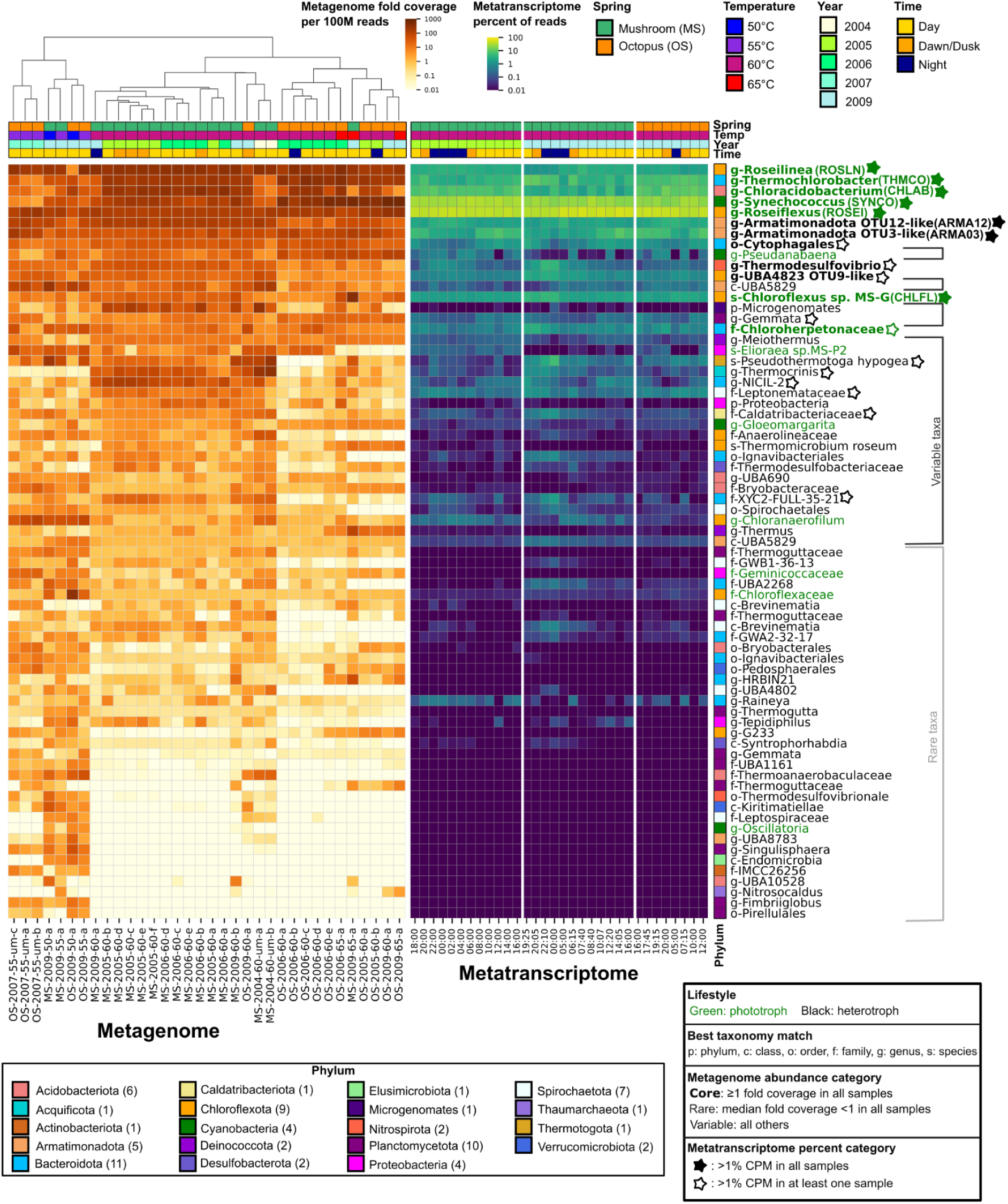
Phototrophic mat microbial community contains core active members, and sample-specific populations. The relative abundance of the 71 genus to species-level taxa groups obtained from the pan-genome ortholog clustering pipeline was ordered by median relative abundance in all 34 metagenome samples. The metagenome samples are ordered by hierarchical clustering on the relative abundance of the 71 taxa (70 bacterial and 1 archaeal). Phyla of the 71 taxa are indicated by colored boxes, with the number of taxa identified for each phylum in parentheses after the phylum name. The taxa are identified by the closest relative in GTDB or by best match to other undermat Mushroom Spring MAGs (see **Methods**). Abbreviations for best taxonomy match names: p: phylum, c: class, o:order, f: family, g: genus, s: species. Metatranscriptome samples are ordered by sample year and time. Sample properties are colored at the top as indicated. Taxa properties: Green text: Phototrophic taxa based on annotations indicating the presence of photosynthesis reaction center proteins, black text: inferred heterotroph based lack of reaction center proteins. **Bold**: Core taxa with at least one fold coverage per 100M reads in all 34 metagenome samples. Rare taxa indicated by light gray brackets are defined as median coverage of less than one-fold per 100M reads in the 34 metagenome samples. Variable taxa indicated by dark gray brackets have median coverage between core and rare cutoffs. Filled stars: the eight taxa with over 1% CPM in all 32 metatranscriptome samples; Open stars: taxa with at least 1% total counts per million in at least one metatranscriptome sample.

**Table 1:** Summary of samples. METAGENOME = designations denote spring, year temp and samples taken at different times during the day for metagenome samples, METATRANSCRIPTOME = sampling series name for metatranscriptome samples. Last row: summary of sampling. Abbreviations: MS = Mushroom Spring, OS = Octopus Spring, letters (a-f) represent individual samples for the metagenome. um=undermat. nd = not done.

| TEMP (°C) | YEAR | SPRING | METAGENOME | METATRANSCRIPTOME |
| --- | --- | --- | --- | --- |
| 60 | 2004 | MS | MS-2004-60-um-a,b | nd |
| 60 | 2005 | MS | MS-2005-60 a, b,c, d, e, f, | MS2005 (12 samples over diel cycle) |
| 60 | 2005 | OS | OS-2005-60 a, b | nd |
| 60 | 2006 | MS | MS-2006-60 a, b,c, d, e, | nd |
| 60-65 | 2006 | OS | OS-2006-60 a,b,c,d,e, 65a | nd |
| 55 | 2007 | OS | OS-2007-55-um-a,b,c | nd |
| 50-65 | 2009 | MS | MS-2009-60a, 50a, 55a, 60b, 65a | MS2009 (12 samples over diel cycle) |
| 50-65 | 2009 | OS | OS-2009-60a, 50b, 55c, 60d, 65e | OS2009 (8 samples over diel cycle) |
| **50- 65 range** | **5 yr range** |  | **34 samples (5 years, 4 temperatures)** | **32 TOTAL (2 years, 1 temperature (60))** |

Sample-specific assembly was performed and the extensive number of samples in this study allowed cross-sampling binning based on variable coverage using MetaBAT 2 ^29^. We identified 1338 metagenome assembled genomes (MAGs) (**Fig. 1, Supplementary Data 2**) (doi:10.6084/m9.figshare.26530315). A common issue with environmental metagenomic datasets is the high fraction of low quality MAGs. To overcome this we applied a pan-genome pipeline with manual curation after automated binning and taxonomic identification. The anvi’o pan-genome pipeline ^30^ was used to identify homologous gene clusters (ortholog groups) between MAGs in the same phylum, and to cluster MAGs into genus- to species-level groups, hereafter referred to as “taxa,” based on the presence of shared ortholog groups in the anvi’o pangenome viewer (**Fig. 1**). The pan-genome pipeline was applied to a single phylum at a time and used the Genome Taxonomy Database toolkit (GTDB-tk) ^31,32^ to identify MAGs. The taxonomic level with the best match in the GTDB was used as the designation for all MAGs belonging to a pan-genome taxon **(Supplementary Data 2**). Only about half of the pan-genome taxa were identified to the genus or species level by GTDB-tk, so to improve resolution of the taxonomic identification, additional curation was performed using fastANI^33^ analysis of MAGs from this study against MAGs derived from a 60°C Mushroom Spring metagenome sample that were not included in GTDB^5,6^ (**Supplementary Data 3)**. We filtered out taxa consisting of a single MAG or very low quality MAGs and 830 MAGs were retained for further analyses. We noted that more than 70% of reads in each sample mapped to the 830 selected MAGs (**Supplementary Fig. 1**), suggesting that these MAGs captured most of the mat phylogenetic diversity.

Of the 830 selected MAGs, 828 MAGs were grouped into 70 taxa from 18 bacterial phyla and 1 taxon in the phylum Thaumarchaeota (2 MAGs were identified as belonging to the archaeal genus *Nitrosocaldus* sp.) (**Fig. 2, Supplementary Data 2**). By comparison, earlier studies from OS and MS reported only eight bins from four Sanger sequenced samples, two of which were completely unidentified, whereas Thiel *et al.* recovered 15 bins from an undermat sample sequenced with Illumina HiSeq^5,6,27^.

We identified a core set of 12 taxa present in all 34 metagenome samples across both springs and all times and temperatures with at least one-fold coverage per 100 million reads (**Fig. 2**, bold names, **Supplementary Data 4**). These included seven phototrophic taxa in four phyla: Acidobacteriota (*Chloracidobacterium*), Chloroflexota (*Chloroflexus, Roseiflexus*, *Roseilinea),* Cyanobacteriota (*Synechococcus*) and Bacteriodota (*Thermochlorobacter,* one in the family Chloroherpetonaceae). Most of these taxa have been previously observed in hot spring samples from this location^2,34^. We use *Synechococcus* to describe the most abundant cyanobacterial genus present in this study, although it has been proposed that it be placed in the genus *Thermostichus* sp. because of its evolutionary distance from the type strain ^13,35^. Five heterotrophic taxa were among the core taxa: Armatimonadota OTU3-like, Armatimonadota OTU12-like, a taxon from order Cytophagales from phylum Bacteroidota, *Thermodesulfovibrio* from phylum Nitrospirota, and a taxon from the Chloroflexota genus UBA4823 related to Thiel *et al.’s* OTU 9^5,6^. Of the heterotrophs, only *Thermodesulfovibrio* spp. has a cultured representative from a different hot spring, and the others except for the Cytophagales taxon have been described only in genomic datasets ^5,6,10,36^. Understanding the role of these heterotrophs in the mat community will require further characterization.

Most of the abundant taxa are found in MS and OS, consistent with previous studies ^2,3^. Both springs are located about 300 meters apart in the Lower Geyser Basin of Yellowstone National Park, and have similar physical and chemical properties, although the source temperature of OS is 90°C compared to around 70°C in MS ^37,38^(**Fig. 1**). However, analysis of data from the 60°C sites at MS and OS, for which we had the most samples (n=22), showed that the Shannon index for alpha diversity performed on the fold coverage data of the 71 taxa was lower in the samples from OS than MS (**Supplementary Fig. 2a**, Kruskal-Wallis statistic=9.232, p-value=0.002, **Supplementary Data 5**). The samples were also significantly different based on the ANOSIM test (R=0.769, p-value=0.001, 999 permutations) on the Bray-Curtis dissimilarity metric for beta diversity (**Supplementary Fig. 2b, Supplementary Data 6**). Together, these metrics suggest that the taxa composition of the phototrophic mats at 60°C in MS and OS are different. This is consistent with a prior study where community composition in the overflow water that passes above the phototrophic mats was reported to be different between OS and MS springs, which was attributed to differences in source temperatures ^39^. We used the ANCOM method ^40^ which identified five taxa that were differentially abundant between the two springs. This includes two taxa from the phylum Spirochaetota, one from the Bacteroidota, a taxon from phylum Chloroflexota, and a genus from phylum Acquificota (*Thermocrinis*) (**Supplementary Data 7, Supplementary Fig. 3**). Of these, only *Thermocrinis sp.* has been described in some detail, as it was previously isolated from the upper outflow channel of OS ^41^. We cannot ascertain whether the differences in community composition are a consequence of chemical, thermal, or biological differences between the springs or a result of historical contingency of colonization. With more intensive sampling of multiple springs^42^ or extensive longitudinal sampling, it may be possible to address the role of community interactions, environmental constraints, and evolutionary history on extremophile microbial communities.

In this study, we focused on the abundant and active taxa in the community, although we identified thirty-five “rare taxa’’ from fourteen phyla which had a median coverage of less than one. In addition there were twenty-four “variable taxa’’ with median coverage values between the “core” and “rare” taxa from fourteen phyla (**Fig. 2**, **Supplementary Data 4**). There are four additional phototrophs in the “variable taxa”, and three in the “rare taxa.” Some of the “variable” and “rare” taxa were present at higher abundances in some samples and we noted that “rare taxa” appear to be specifically found at lower temperatures (50°C and 55°C) or in undermat samples. However we did not perform further significance testing. Some rare taxa appear to be specific to each spring at 60°C and are discussed above. Samples from lower temperatures and the undermat layer had more diverse communities, as indicated by higher Shannon indices (**Supplementary Data 5**), which is consistent with previous results ^5,43,44^.

*Transcriptional activity measured at 60*°*C is primarily derived from a few highly active taxa*

To identify taxa that contributed most to community activity, we carried out metatranscriptomic analysis of samples collected every 2-5 hours over a 24 hour period at a 60°C site. These samples were taken at three locations and times: MS in late June-early July, 2005 (MS2005) and in July 2009 (MS2009) and OS in July 2009 (OS2009) (**Fig. 1**, **Table 1, Supplementary Data 1**). Ribosomal-depleted RNA reads were passed through a quality control filter using a standard pipeline (**Fig. 1**, **Table 1, Supplementary Data 1**) and then aligned to a reference containing the selected 830 MAGs (doi:10.6084/m9.figshare.26530315). To assess biological activity of genes, only reads within annotated coding sequences (CDS) were counted (10-30% of aligned reads per sample)(**Supplementary Data 1**)(doi:10.6084/m9.figshare.26530315). We aggregated the RNA counts of MAGs for each pan-genome taxon by summing counts from the ortholog groups identified by anvi’o (hereafter referred to as genes) (**Supplementary Data 8**). The counts were normalized for each time series to counts per million (CPM) (**Supplementary Data 9**).

We do not expect organism relative abundance to have a major impact on the interpretation of activity because metagenome samples taken within 24 hours of each other tended to have very high similarity compared to other samples, as measured by Bray-Curtis beta diversity, suggesting limited change in relative abundance over short time scales (**Supplementary Data 6**). We detected RNA for 0-95% of the predicted CDS per genus (**Supplementary Data 10**), with those at the lower end reflecting the range of relative abundances based on DNA analyses in the 60°C mat samples.

We focused on “active taxa’’, i.e.taxa present at the level of at least one percent or higher of the total number of aligned reads in all 32 metatranscriptome samples (varied from 1.2%-55.6%) (**Fig. 2**, filled stars). Of these, six taxa were identified as phototrophs (*Roseiflexus* (ROSEI)*, Chloracidobacterium* (CHLAB)*, Thermochlorobacter* (THMCO)*, Roseilinea* (ROSLN)*, Chloroflexus* (CHLFL), and *Synechococcus* (SYNCO)) and two were heterotrophs in phylum Armatimonadota (Armatimonadota OTU3-like (ARMA03) and Armatimonadota OTU12-like (ARMA12)). For simplicity, we use the short abbreviations mentioned in parentheses in some figures. This “first tier” of the most active taxa (**Fig. 2**, filled stars) is consistent with the most abundant taxa (**Fig. 2**, bold names). A “second tier” of eleven additional taxa made up at least 1% of total aligned and counted RNA reads at one or more time points (**Fig. 2**, open stars), but due to low counts in some samples, their gene expression patterns were not specifically analyzed further.

Typically, metatranscriptomic analyses are based on *de novo* assembly of transcripts that are annotated using a well-curated database^45,46^. A second approach maps the RNA reads to available reference genomes, selected relevant MAGs, or a reference database of sequences ^47–51^. Because of the paucity of high-quality reference genomes for these mat communities, assigning assembled transcripts to a meaningful taxonomic level or obtaining high gene coverage by mapped reads to reference genomes was challenging. So, we mapped reads to all MAGs and summed the counts per ortholog group for each taxon identified in the pan-genome to overcome the problem that some highly abundant, active taxa did not have reference genomes, had fragmented MAGs (**Supplementary Data 2**), or had closely related genomes that were not easily separated into unique MAGs ^5,18,25,52^. This approach allowed us to capture a higher fraction of the reads compared to mapping them to dereplicated MAGs or reference genomes (data not shown) while still capturing a genus-level context for the genes.

### Active taxa exhibit peaks of gene expression at different times of the diel cycle

Initially, we sampled some of the highly expressed genes (defined as mean CPM greater than one) in the eight most active taxa to identify diel regulation of genes. Approximately 10% of all genes were highly expressed (24,906 genes out of the 233,893 annotated genes). Highly expressed genes were detected in 50 of the 71 taxa (**Supplementary Data 10**). In the eight most active taxa, between 34% and 82% of genes per taxon were highly expressed (**Supplementary Data 10**).

To get an overview of gene expression patterns in each of the active taxa, we visualized the expression patterns of a subset of highly expressed genes over the diel cycle in heatmaps in **Figure 3a**, which shows that each of the eight taxa appear to have distinct peak expression patterns. As in **Figure 1**, we qualitatively describe the time of day by the lower color bar with yellow being day, orange representing dawn or dusk, and blue for night as indicated in **Supplementary Data 1**. Most gene expression patterns varied as a function of the diel cycle in both the phototrophs and heterotrophs. There were few genes with constitutive expression patterns over the diel cycle, which would be visualized as the same color across all time points.

**Figure 3:**
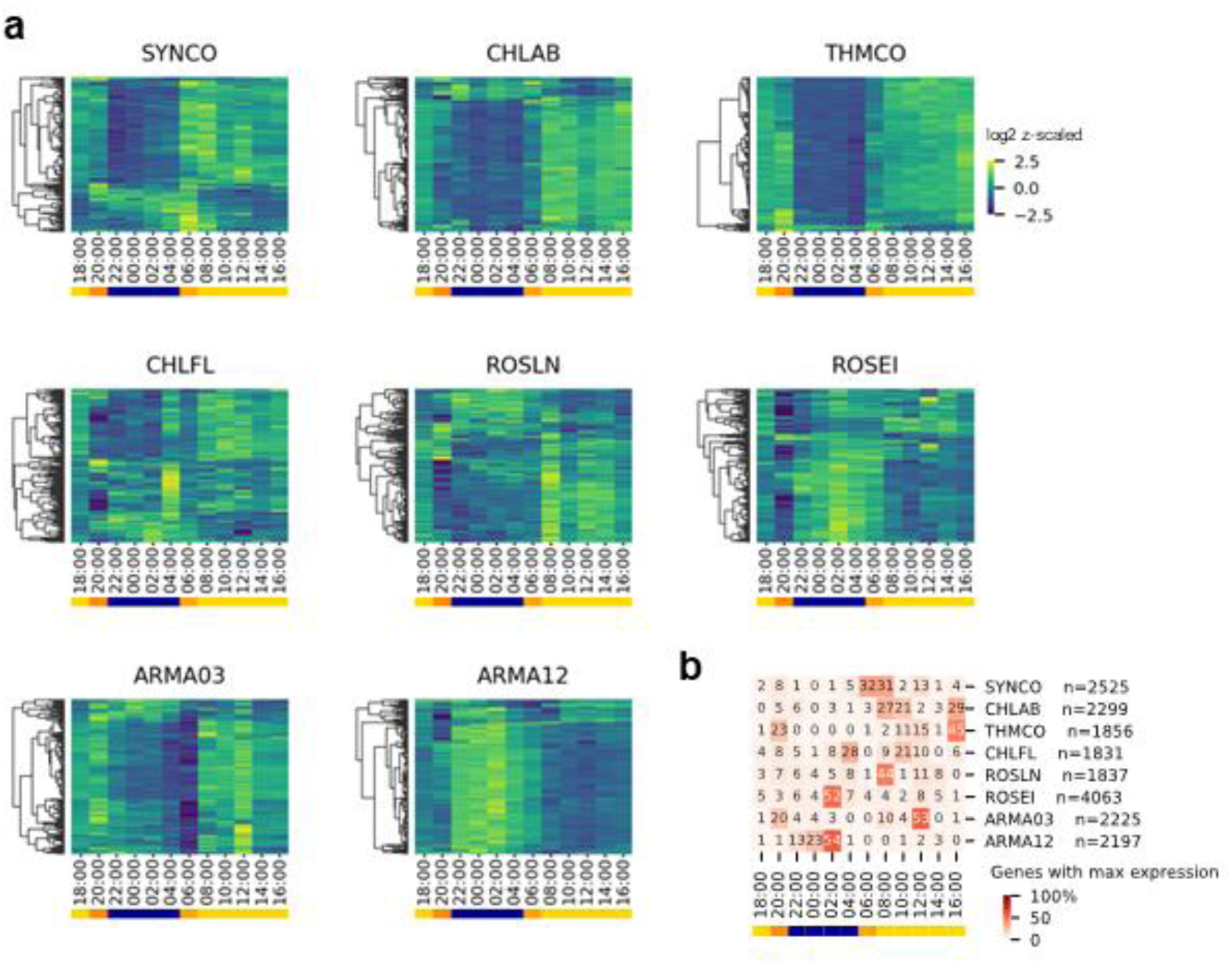
Gene expression of the eight most active taxa varies during the diel cycle. **a.** Log2 normalized and z-scaled expression in MS2005 of 500 randomly selected highly expressed (mean CPM >1) genes of the eight most active taxa. Sampling of a subset of genes was performed to select genes from across the genome and allow visualization within the limits of the figure. Dendrograms were generated using seaborn clustermap with a correlation distance matrix and complete clustering. **b**. Heatmap of percent of highly expressed gene clusters per taxon that had maximum expression at each time point. n= number of genes analyzed per taxon. Bars at the bottom indicate qualitative time of day: yellow: day, orange: dusk or dawn, blue: night as described in **Supplementary Data 1**.

To tabulate the differences in expression patterns among the taxa, we summarized by counting the number of genes per taxon with the maximum expression values at each time point (**Fig. 3b, Supplementary Fig. 4**). Six of the eight most active taxa (*Synechococcus*, *Chloracidobacterium*, *Thermochlorobacter*, *Chloroflexus*, *Roseilinea*, and Armatimonadota OTU3-like) had >50% of their genes peak during the day (yellow bar at bottom of **Fig. 3b**). On the other hand, Armatimonadota OTU12-like and *Roseiflexus*, had >50% of their genes peaking at night (blue bar at bottom of **Fig. 3b**). Among the six taxa that had a majority of day-peaking genes, we noticed a further distinction. Armatimonadota OTU3-like and *Thermochlorobacter* had 20% or more of their genes which exhibited peak expression during dusk, while in *Synechococcus,* 20% or more genes exhibited peak expression at dawn (orange bar at bottom of **Fig. 3b**), suggesting particular transcriptional regimes that are more responsive to the transition periods of dusk or dawn. An example of dawn-responsive gene expression in the mat are genes encoding nitrogenase, which have a minor peak at night and a second larger peak at dawn^23^ consistent with nitrogenase activity that peaks in the early morning. We noticed that 53% of Armatimonadota OTU3-like genes had their peak expression at noon (light period), while in Armatimonadota OTU12-like, 54% genes had peak expression at 02:00 h (dark period). This striking contrast in peak activity of the majority of genes suggests that these two taxa may have significantly different metabolic lifestyles, which we describe in more detail in the section on Armatimonadota.

### Response to diel cycle in sentinel pathways

The previous analyses suggested that the eight most active taxa could be broadly categorized into either “day-active” or “night-active” taxa. To validate and extend this distinction, we examined expression patterns of a few sentinel genes/pathways known or expected to be regulated by light, anoxia or circadian rhythms by selecting marker annotations using KEGG Orthology (KO)^53^ and Cluster of Orthologous Groups (COGs)^54^, see **Methods**) ^9,20–23,55,56^ It is visualized for MS2005 in **Figure 4**. We show the MS2009 data in **Supplementary Figure 5a**, and OS2009 in **Supplementary Figure 5b**, while the underlying data is in **Supplementary Data 11**.

**Figure 4:**
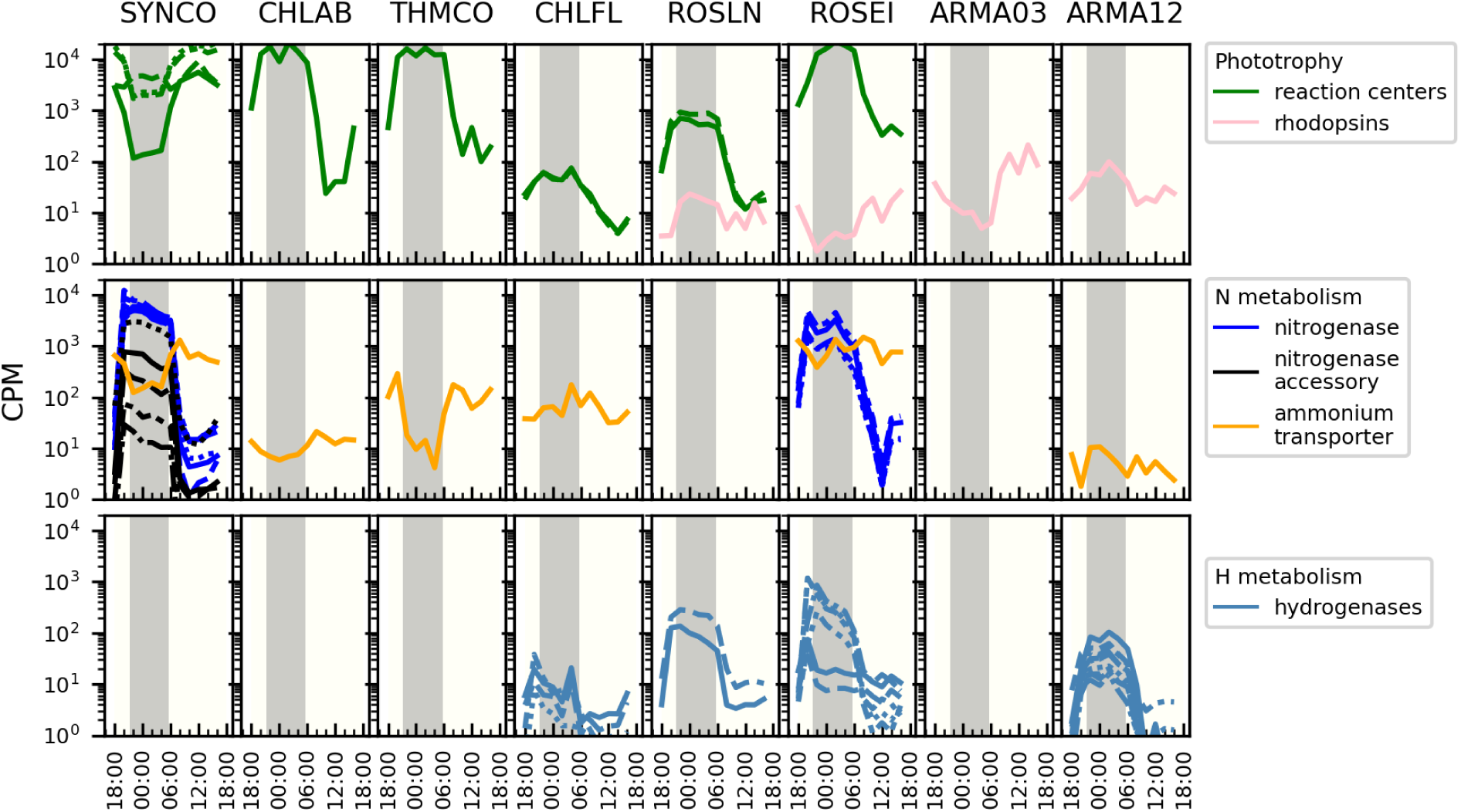
Response of sentinel pathways to the diel cycle in the eight most active taxa. Gene expression from the MS2005 metatranscriptome is plotted for the eight most active taxa. Panels without curves indicate that a gene containing that annotation was not expressed in that taxon. Each line per panel is a separate gene. Gray bar: Night time. CPM: counts per million. Underlying data is found in **Supplementary Data 11.**

#### Phototrophy

As expected, *Synechococcus* had higher expression of genes encoding photosynthetic reaction center proteins during the day^9,20,23,56^ while in the anoxygenic phototrophs, reaction center gene expression was higher at night ^20–22^. In addition, we observed rhodopsin expression in four of the eight active taxa which might be indicative of an alternative to (bacterio)chlorophyll-based phototrophy (**Fig. 4**, **Fig. 2**, names in green). Both Armatimonadota taxa also expressed rhodopsins although the expression was not specifically responsive to the diel cycle. Rhodopsin expression was higher at night in *Roseilinea*, but higher during the day in *Roseiflexus* (**Fig. 4**, **Supplementary Fig. 5**). This suggests that there are inputs into the regulation of rhodopsins in addition to light ^57,58^ but further experimentation will be required to substantiate these findings.

#### Nitrogen fixation and assimilation

Nitrogenase activity in the unicellular *Synechococcus* has been shown to have peak activity in the early morning consistent with the model that nitrogenase requires high levels of ATP and reductant which are readily available in the early morning hours when photosynthesis is increasing. However, later in the day the increasing level of oxygen inactivates the nitrogenase. On the other hand, peak expression of the *nifHDK g*enes occurs at night^22,23^. Our metatranscriptomic data confirms and extends these results by showing that the major *nifHDK* genes encoding the nitrogenase enzyme as well as the nitrogenase accessory subunit expression follow this trend. Nitrogenase accessory genes were not encoded or expressed in *Roseiflexus* MAGs, consistent with previous genome analyses and reports of its inability to grow on N_2_ ^6,59^(**Fig. 4, Supplementary Fig. 5**). We observed expression of ammonium transporters (Amt) in six of the eight taxa. This is consistent with the observation that there is very little fixed nitrogen in the source water, so organisms capable of nitrogen fixation in the mat or release of nitrogen compounds by cell lysis are the major sources of fixed nitrogen ^38,55^.

#### Hydrogenases

H_2_ concentration increases in the mats at night with a second peak at dawn, possibly concurrent with nitrogenase activity, since hydrogen is a known byproduct of nitrogenase activity^8^. Consistent with this, hydrogenases in Armatimonadota OTU12-like, *Chloroflexus*, and *Roseiflexus* exhibited elevated levels of expression at night when the mats are anoxic and amenable to enzyme assembly (**Fig. 4, Supplementary Fig. 5)**. Notably, *Synechococcus* appears to lack hydrogenases, suggesting that hydrogen, which is a byproduct of fermentation and nitrogen fixation activity by this organism, could serve as a source of reductant/energy for other hydrogenase-containing organisms in the mat. To test this hypothesis, additional follow-up experiments will be required.

### Consensus module detection reveals co-expression of genes within and between taxa

Next, we wished to extend this analysis beyond known sentinel genes and pathways to discover biological activities or interlinked behaviors in these coexisting microbes. To do so, we identified groups of genes with similar expression patterns across the entire community using weighted gene correlation network analysis (WGCNA) with consensus module detection on the highly expressed genes of the metatranscriptomes ^60,61^. Briefly, this method takes advantage of the three metatranscriptome time series to independently calculate the correlation between genes in each dataset. Subsequently, the method discovers genes that are consistently co-expressed in all three time series and uses hierarchical clustering to identify clusters of genes that are co-expressed (or modules). There are only a few reports of this method being used to dissect microbial community interactions ^62^ but if there are appropriate replicate datasets, it can be a powerful discovery tool.

Using this method, most of the highly expressed genes (24,701 of 24,906 genes) were placed into fifty-four modules (ME1-54). The few remaining genes (205, labeled ME0) did not exhibit consistent co-expression across all three time series and could not be placed in a specific module (**Fig. 5**, **Supplementary Data 12**). By performing simultaneous WGCNA on all organisms, we were able to infer co-expression between taxa (i.e. genes in the same WGCNA module indicates high correlation). Expression patterns of genes in each WGCNA module were summarized by the module eigengene, the first principal component of each module, as a representative expression pattern. We used the eigengenes to describe the relatedness of modules through hierarchical clustering of their expression patterns across all three time series (**Fig. 5b**, **Supplementary Fig. 7, Supplementary Data 13, 14**). To simplify comparisons, we categorized WGCNA modules into four groups (morning/day-, night/morning-, evening/night-peaking, or variable patterns between time series) (**Fig. 5b**) based on peak expression time relative to measured light intensity (**Supplementary Data 1**).

**Figure 5:**
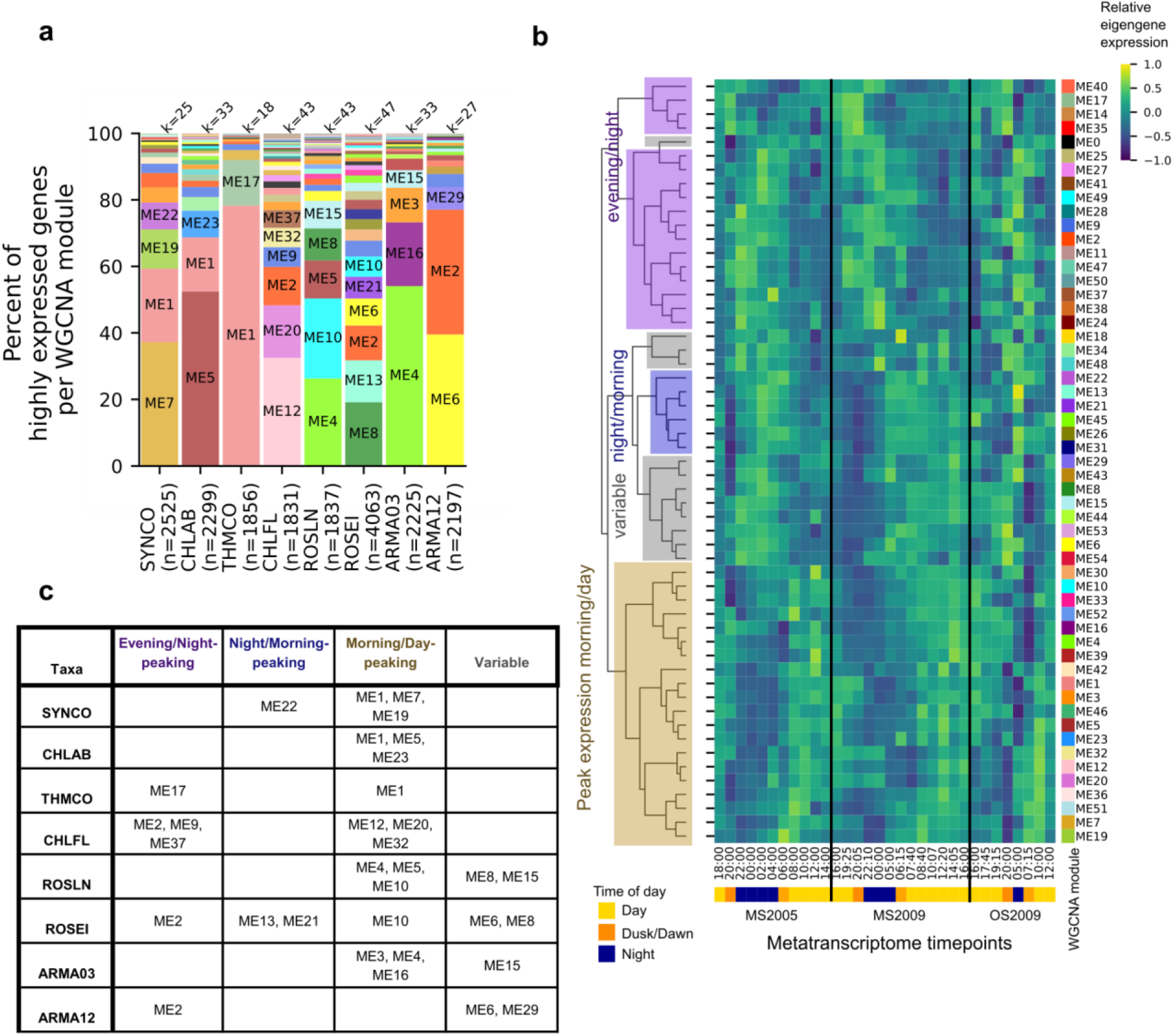
Consensus WGCNA modules in show differences and similarities in expression patterns between taxa. **a.** Percent of highly expressed genes per WGCNA module in active taxa. Modules containing 5% or more of highly expressed genes per taxon are labeled. ME0 (black fill) represents genes that did not fit in any of the numbered consensus modules. n=number of highly expressed genes per taxon, k=number of WGCNA modules per taxon. Raw gene counts are plotted in **Supplementary Fig. 6. b**. Eigengene expression for the 54 modules and ME0 across all three sampling series. Underlying data is found in **Supplementary Data 13.** Colors on the right correspond to the same colors in (**a**). Labels were placed on the dendrogram describing the relationship between eigengenes. Except for the “variable” category, eigengenes had qualitatively similar peak expression times in each of the three time series (MS2005, MS2009, and OS2009). The Pearson correlation between eigengenes (**Supplementary Fig. 7, Supplementary Data 14**) was used for the dendrogram linkage. **c**. Modules containing 5% or more of highly expressed genes per taxon organized by eigengene peak expression. Colors match the eigengene categories on the dendrogram in **(b)**.

Using this pipeline, we made a number of unexpected observations. First, we found that most WGCNA modules tend to be dominated by a single taxon. More than 50% of the genes in 47 of the 54 modules were from a single taxon. In 16 modules, 90% of the genes were from a single taxon (**Supplementary Fig. 8**). Thus, we might initially hypothesize that gene expression is not very strongly coordinated *between* taxa but primarily *within* taxa or in other words, it is “organism specific”. However we did find examples where six modules contained genes from more than 20 genera (**Supplementary Fig. 8**). This included the largest WGCNA module, ME1, which is a morning/day-peaking module (n=2682 genes, 21 taxa) and ME2 which is an evening/night-peaking module (n=2521 genes, 40 taxa). This result suggests that there is coordination or convergence in expression patterns across taxa. The fact that we did not find extensive coordination by strict module classification is not unexpected as this method is sensitive to the relative expression and the magnitude of the change in gene expression, which varies widely among taxa (**Supplementary Data 15**).

While we observed limited co-expression in many modules based on inclusion of multiple genera in each module, we did find some similarity in expression patterns when modules were hierarchically clustered and categorized by peak expression timing (**Fig. 5b, Supplementary Fig. 7**). Notably, the two Armatimondota taxa, *Chloracidobacterium*, *Thermochlorobacter* and *Synechococcus* had 38-78% of their highly expressed gene clusters in a single module (**Fig. 5a, c, Supplementary Fig. 6**), and all these modules were morning/day-peaking. The striking exception was Armatimonadota OTU12-like which had many of its highly expressed genes in an evening/night module (ME2). In contrast, the three Chloroflexota exhibited more complex partitioning of their gene expression patterns, and their module sizes were generally smaller than the other taxa. The largest modules in *Chloroflexus* and *Roseilinea* were morning/day-peaking, whereas those in *Roseiflexus* had no consistent expression patterns across all three time series or were night-peaking (**Fig. 5**, **Supplementary Fig. 6**). These results of peak expression timing were consistent with the observations shown in **Fig. 3**.

### Overrepresentation of biological functions in WGCNA modules reveals similarities in expression timing

To further dissect the variation in expression patterns within and between taxa we linked WGCNA modules to biological functions using Clusters of Orthologous Genes (COG) Categories ^54^. To determine if a biological function was significantly overrepresented in the WGCNA module of each taxon, we performed hypergeometric overrepresentation analysis (ORA) against the total number of highly expressed genes annotated with COG categories in that taxon.

We found 89 instances of COG categories in the active taxa that were significantly overrepresented in modules (**Fig. 6**, **Supplementary Data 16**). For instance, the translation and ribosome structure and biogenesis category (Category J) was significantly overrepresented in morning/day-peaking modules in all six phototrophs and Armatimonadota OTU3-like (**Fig. 6**).

**Figure 6:**
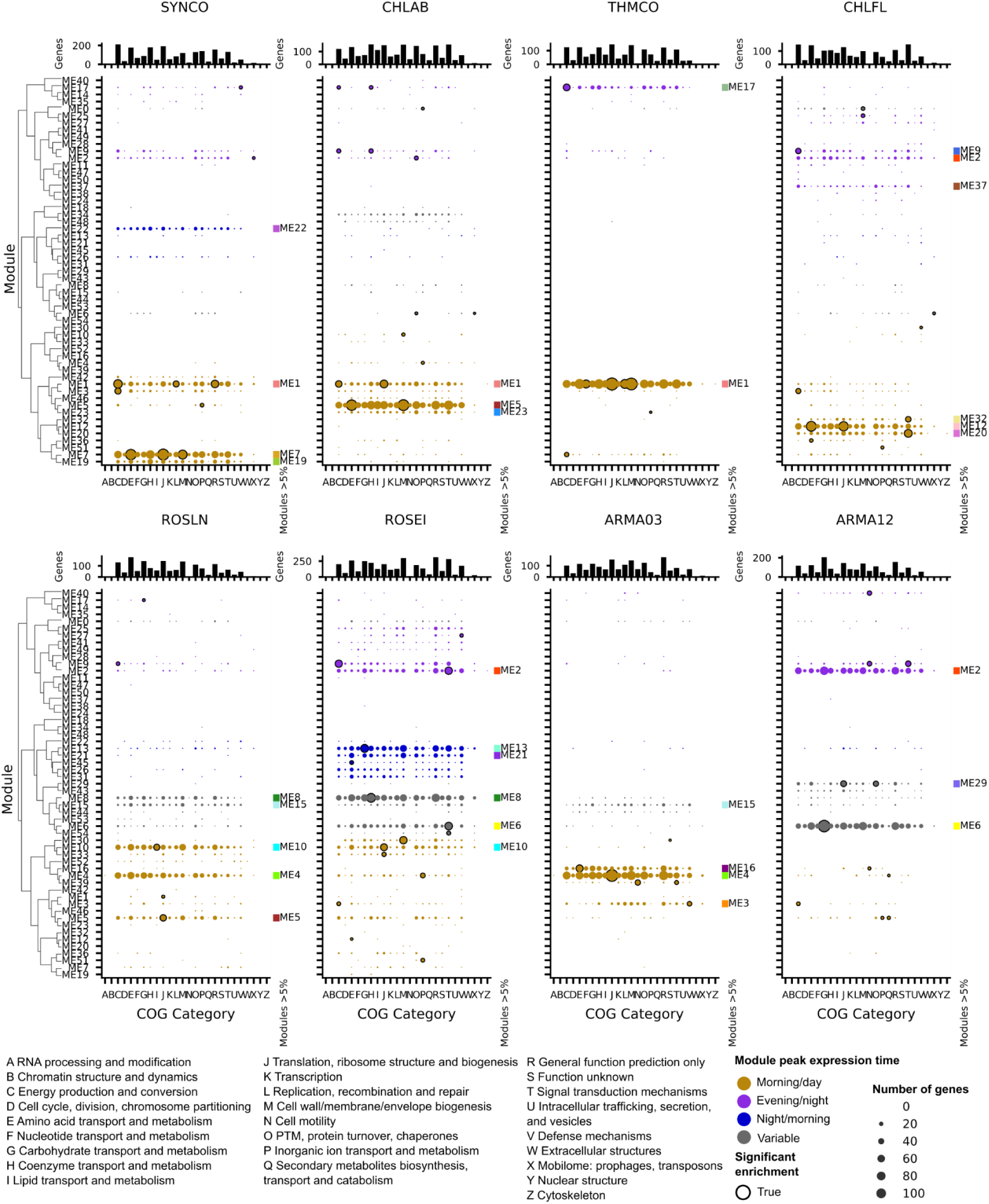
COG Category overrepresentation per WGCNA module in active taxa. For each of the eight most active taxa the number of genes per WGCNA module is plotted per COG Category. Circle size represents the number of genes. Significant overrepresentation of a COG category in a module is indicated by a black outline. Module ordering and coloring is the same as in Figure 5b based on the hierarchical clustering of module eigengenes. The top bar chart for each taxon is the total number of genes per COG category in that taxon. Modules representing 5% or more of the highly expressed genes per taxon are highlighted to the right, colors match Figure 5ab. Associated data is found in **Supplementary Data 16**.

Selected example genes are plotted in **Fig. 7** for the MS2005 dataset and in **Supplementary Fig. 9** for the MS2009 and OS2009 datasets **(Supplementary Data 17).** This suggests that these seven taxa are likely to have high rates of protein synthesis in the morning and day. We hypothesize that as photosynthesis peaks during the day, it provides the energy required for energetically intensive cellular processes such protein synthesis and ribosome biogenesis.

**Figure 7:**
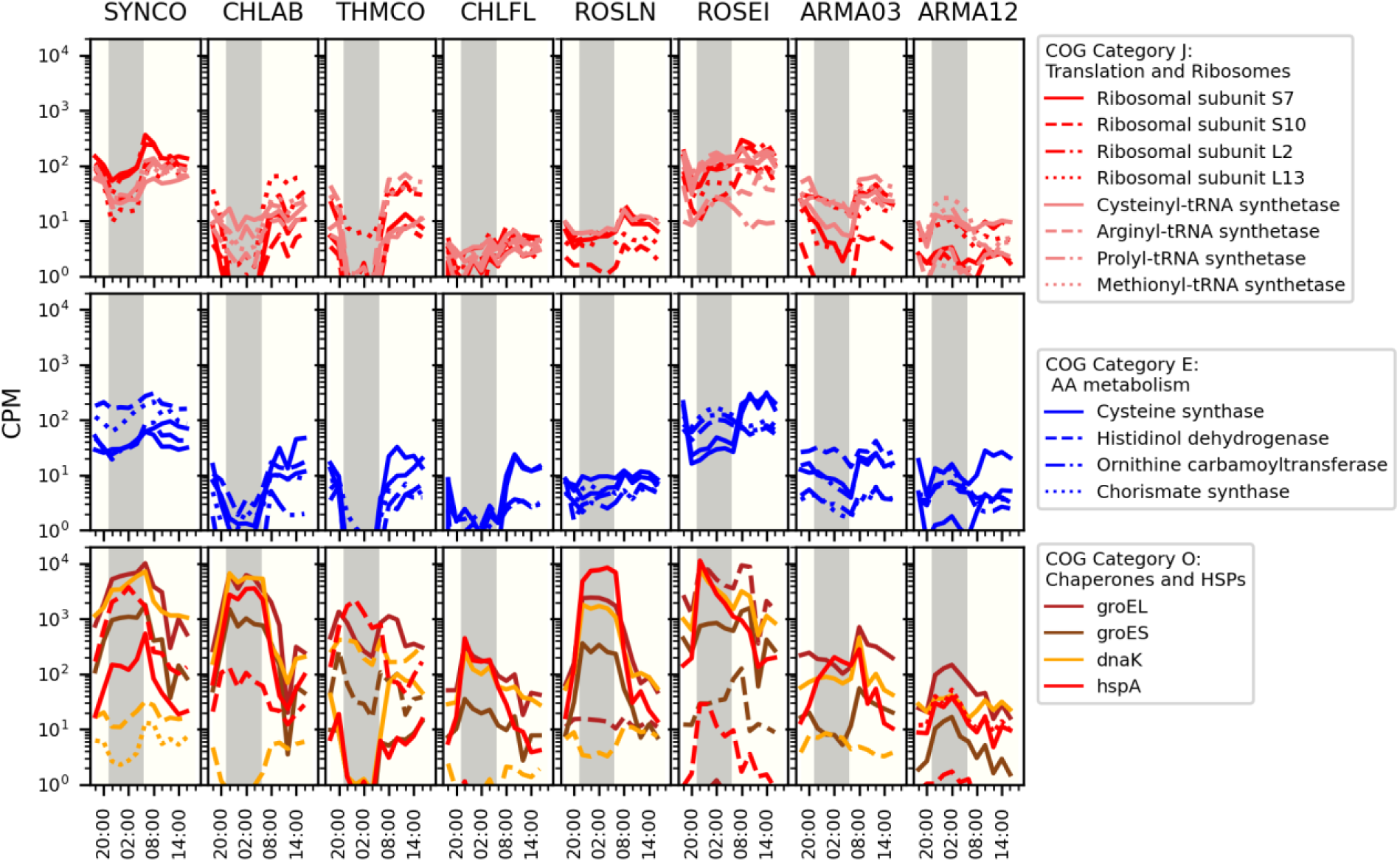
Expression patterns of selected genes from the COG Category overrepresentation analysis. A subset of genes from COG Categories of interest were selected to be visualized for the eight most active taxa using the MS2005 metatranscriptome. Each line is a different gene. Top: Category J, Translation and Ribosomes. Middle: Category E, Amino acid (AA) metabolism. Bottom: Category O, Chaperones and heat shock proteins (HSPs). CPM: counts per million. Underlying data is found in **Supplementary Data 17**.

While this timing of peak ribosome subunit expression is consistent with peak gene expression in other pathways in most of these seven taxa, many genes in *Roseiflexus* are highly expressed at night. One possible explanation for some pathways is that translation is temporally decoupled from transcription and occurs when more ribosomes are present. Coupled transcriptional and proteomic data would be required for validation but to our knowledge, very few studies have been designed to assess coupling of transcription, translation and function in environmental communities^9,23^. Klatt *et al*. noted that there was a temporal discrepancy in *Roseiflexus* between specific transcriptional profiles and when a measured function, such as light-driven CO_2_ fixation was highest based on *in situ* data. They suggested it may be due to post-translational regulation,^22^ although our data suggest an alternative scenario. Interestingly, the one notable exception to this pattern of morning ribosome expression is Armatimonadota OTU12-like, which is consistent with the finding that it appears to have the majority of its activity at night (**Fig. 3, 4, 5, 6**).

Another example of conserved peak expression timing among taxa was that amino acid biosynthesis (Category E) was significantly overrepresented in morning or day peaking modules in Armatimonadota OTU3-like, *Chloroflexus, Roseiflexus* and *Synechococcus*, and a subset of specific genes in these pathways exhibit dawn peaks in these taxa (**Fig. 6, 7, Supplementary Fig. 9, Supplementary Data 16, 17**). This is consistent with observations that peak amino acid biosynthesis in the mat occurs in the late morning, based on *in situ* metabolomics^28^. Cell wall biosynthesis genes (category M) were significantly overrepresented in morning/day-peaking modules in *Roseiflexus* (ME30) and *Synechococcus* (ME7), also suggesting that these two taxa have similar timings of cell growth and division (**Fig. 6**). Therefore, despite differences in pigmentation and sources of fixed carbon and reductant ^2^, many of the phototrophs have increases in expression of genes associated with cell growth and division in the morning, suggesting some convergence in timing of the cell cycle. Additional *in situ* data types such as microscopy to count dividing cells, measuring protein synthesis rates, or laboratory experiments monitoring the cell cycle will be required to strengthen this hypothesis.

We also used the ORA approach to discover pathways that are active in the absence of light energy in the phototrophs. Some energy production and conversion (Category C) genes of the anoxygenic phototrophs *Thermochlorobacter* (ME17), three Chloroflexota and *Chloracidobacterium* (ME9) were significantly overrepresented in night-peaking modules (**Fig. 6, Supplementary Data 16**). These genes encoded hydrogenases, in those organisms that have them (**Fig. 4, 6**), and genes/proteins expected to be more highly expressed at night, including quinol oxidases, NADH oxidoreductases, ferredoxins and flavodoxins.

The ORA approach also detected genes encoding chaperones from category O (post-translational modification, chaperones and heat shock proteins) that were significantly overrepresented in a night-peaking module (ME2) in *Chloracidobacterium* (**Fig. 6, Supplementary Data 16**). The occurrence of a night-specific chaperone response prompted us to determine if this chaperone expression pattern was unique to *Chloracidobacterium*, or a more general phenomenon. We examined expression patterns of genes encoding specific chaperones across the other seven active taxa (**Fig. 7**, **Supplementary Fig. 9, Supplementary Data 17**). The four other anoxygenic phototrophs exhibited a relative increase in expression of some of these chaperones at night. In contrast, *Synechococcus* chaperones tended to peak around dawn. Some taxa had multiple *dnaK* orthologs in the genus-level pan-genomes; each copy exhibited opposite expression patterns, suggesting different roles for each copy **(Fig. 7, Supplementary Fig. 9, Supplementary Data 17**). We speculate that the timing of expression may be synchronized to peak at specific times for a number of reasons which include (i) control processes that are particularly susceptible to light or oxygen, (ii) limited to a specific phase of the cell cycle, as observed in *Caulobacter crescentus,* (iii) and/or are controlled by the circadian clock in cyanobacteria ^63,64^. Currently, we do not have enough data to distinguish between these alternative explanations.

### Abundant Armatimonadota have different predicted carbon sources and timing of expression

Several of the active and abundant phototrophic taxa have been described ^15,17,21,22,24,56^, whereas the role of heterotrophs in the alkaline hot spring phototrophic mats is underexplored ^10,65^. We detected MAGs from five genera from three classes in the phylum Armatimonadota. All are present in both springs at multiple temperatures. Armatimonadota OTU3-like from class Fimbirimonadia and Armatimonadota OTU12-like from class HRBIN16 were among the most active genera at 60°C and have been previously reported in MS 60°C undermat metagenome samples ^5,6^ (**Fig. 2, Supplementary Data 2, Supplementary Fig. 10**). We hypothesized that their starkly different expression patterns [either day-peaking (Armatimonadota OTU3-like) or night-peaking (Armatimonadota OTU12-like)] may be linked to their genomic and metabolic potential, so we carried out additional manual curation on their MAGs with a focus on their carbon and energy utilization pathways (see **Methods**) (**Fig. 8, Supplementary Data 18, 19, 20, 21, 22)**.

**Figure 8:**
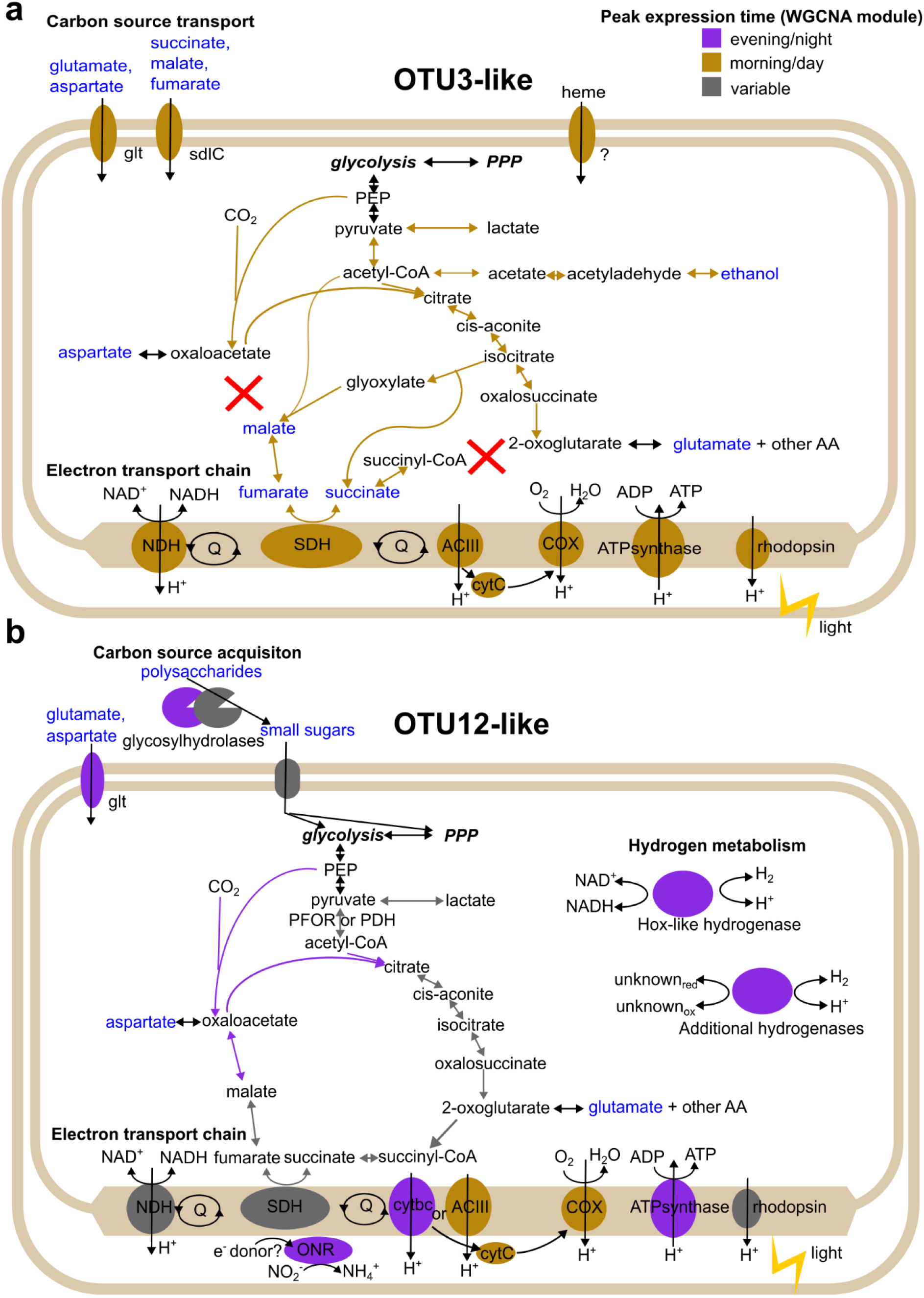
Expression times of major metabolic genes in Armatimonadota OTU3-like and OTU12-like. **(a)** Armatimonadota OTU3-like, **(b)** Armatimonadota OTU12-like. Colors of enzymes are based on their WGCNA module and eigengene expression time (Fig. 5). Metabolites in blue text were predicted to be used by the GapMind webserver. Abbreviations: PPP: pentose phosphate pathway, PEP: phosphoenolpyruvate, AA: amino acids, NDH: NADH dehydrogenase, SDH: succinate dehydrogenase, Q: quinone pool, ACIII: alternative complex III, COX: cytochrome c oxidase, cytC: cytochrome c, cytbc: cytochrome bc, ONR: putative octoheme nitrite reductase, glt: glutamate transporter, sdlC: dicarboxylate transporter.

Armatimonadota OTU3-like has the capacity to use fatty acids and certain amino acids as carbon sources (**Fig. 8a** blue text**, Supplementary Data 18).** Like in *Fimbriimonas ginsengisoli*^66^, a better characterized member of this class, there is an incomplete TCA pathway lacking malate dehydrogenase and oxoglutarate dehydrogenase, but it has glyoxylate shunt enzymes malate synthase and isocitrate dehydrogenase, suggesting an alternative flux in this organism compared to the canonical TCA cycle. While Armatimonadota OTU3-like has an aerobic respiratory pathway with an alternative complex III, it lacks heme biosynthesis, suggesting that it must acquire it from the environment. Consistent with this, subunits of candidate heme transporters were identified and expressed *in situ* (**Supplementary Data 21**). We did not detect any pathway for anaerobic respiration, but it may be able to ferment via the activity of a lactate dehydrogenase, an aldehyde dehydrogenase or an alcohol dehydrogenase. We observed expression of a rhodopsin gene (**Fig 4**), suggesting that Armatimonadota OTU3-like may be able to supplement its energy production/generation of a proton motive force using light (**Fig. 8a**). Many of these pathways are upregulated during the day (**Fig. 3, 4, 5, 6, 8a**), suggesting that its preferred carbon sources are more bioavailable during the day. Consistent with this observation, whole mat polar metabolomics suggested an increase in the relative abundance of some organic acids which Armatimonadota OTU3-like may be able to use in the afternoon^28^.

In contrast, Armatimonadota OTU12-like is predicted to use small sugars including sucrose, cellobiose, glucose, and xylose (**Fig. 8b** blue text**, Supplementary Data 18**). Carbohydrate metabolism (COG category G) was overrepresented in ME6, highly expressed in the evening, night or early morning in the three metatranscriptome series (**Fig. 5, 6**). More secreted glycosylhydrolases per MAG were predicted by dbCAN2 in Armatimonadota OTU12-like than Armatimonadota OTU3-like (mean=16.5 vs 5 per MAG), suggesting that Armatimonadota OTU12-like can degrade extracellular polysaccharides (**Supplementary Data 19**). Whether these carbohydrates are found in the extracellular polymeric substances (EPS) of the mat is not yet known. Unlike Armatimonadota OTU3-like, Armatimonadota OTU12-like contains a complete TCA cycle, multiple bidirectional hydrogenases, a candidate octoheme nitrite reductase, and multiheme cytochromes of unknown function that are predicted to be membrane-embedded (**Supplementary Data 20, 21, 22**). This suggests that there may be alternative electron flow pathways that switch between anaerobic respiration at night and aerobic respiration in the day (**Fig. 8b**). Like Armatmonadota OTU3-like, Armatimonadota OTU12-like also harbors an expressed rhodopsin gene (**Fig. 4, 8a**). A lactate dehydrogenase was also identified, suggesting the potential for fermentation. Many genes are upregulated at night (**Fig. 3, 4, 5, 6, 8b**), although how timing of gene expression relates to the local availability of saccharides is not known. These results suggest differential diel activities and metabolisms of the Armatimondota in hot spring phototrophic mats, which was only previously inferred by genome sequences ^5,6,67^ and suggest possible enrichment strategies to obtain these organisms for which no type strain is currently available.

Here we present a detailed census of microbial taxa present in OS and MS and identify twelve taxa as the core members of the MS and OS mat community. However, relative abundances suggest differences in community compositions between each spring and further support the conclusion that somewhat different communities exist at different temperatures. We analyzed the expression patterns of the most active taxa at 60°C and highlighted possible interactions between taxa. Using the metatranscriptome, we identified eight taxa which are the most transcriptionally active members of the community and notably, all eight had gene expression patterns that correlated with the diel cycle. Our analysis revealed some similarities in expression patterns for pathways and functions, but also highlights major differences in expression patterns such as in the two Armatimonadota heterotrophs.

This sets the stage for understanding community level interactions but we acknowledge some caveats about our datasets and current conclusions. For instance, we do not know to what extent diel variation in transcript abundances is reflected at the protein level ^55,68,69^. Temporal shotgun metaproteomics of the mat will help determine this; so far, such analyses have only been performed for single time points^16,70^. We know that EPS is important for mat structure and may be a source of organic carbon for some mat organisms, as has been observed in other types of microbial mats ^71–73^. While whole mat shotgun polar small molecule metabolomics has been performed ^28^, the dynamics of extracellular polymeric substances (EPS) composition and abundance has not been characterized. We have evidence of metabolite transfer between cyanobacteria and filamentous cells from early radio-labeling experiments^74,75^, however updated techniques will be better able to resolve the identity of both the organisms and metabolites involved. These include activity measurements *in situ*, such as SIP-metagenomics with different labeled carbon isotopes^76^ or SIP-metaproteomics^77^. Our study did not specifically address community abundance, dynamics and activity as a function of mat depth and diel fluctuation in light or oxygen levels which would provide a fine-grained understanding of spatiotemporal mat dynamics.

## Methods

### Sample sites and collection

Mat cores were collected at Mushroom Spring (44.53861N, 110.79791W) and Octopus Spring (44.53401N, 110.79781W) on the dates and times provided in **Table 1 and Supplementary Data 1.** Cork borers (∼0.5 cm to 1 cm) were used to collect mat cores, which were immediately frozen in liquid nitrogen until later use. Irradiance at the top of the mat was measured June 30-July 1, 2005, July 28-30, 2009, and October 1, 2005, as described in Jensen *et al.* ^9^ (**Fig. 1, Supplementary Data 1**). For the MS2005 metatranscriptome time series, a sample taken at 12:00 PM from October 1, 2005 was used to replace the original July 1, 2005 12:00 PM sample.

### DNA and RNA extraction

A modified phenol:chloroform:isoamyl alcohol extraction was performed on mat core samples to extract total DNA and RNA^55^. ReadyLyse (LUCIGEN) and lysozyme were added after bead-beating. For DNA extraction, RNase A (NEB) was additionally added. Following phenol-chloroform extraction, the nucleic acids were precipitated overnight in 100% ethanol. For the RNA samples, RNase A was omitted following lysis. Following precipitation of total nucleic acids in ethanol, superase (Thermo Fisher) was added at ∼0.5 U /uL in pH 8.0 TE for resuspension, followed by DNase (NEB) digestion for 10 min in 100 uL with 2 U/uL superase, and then immediately purified with Qiagen RNeasy Mini-elute for total RNA. DNA and RNA were quantified with Qubit (ThermoFischer) and Fragment Analyzer (Agilent).

### Library preparation, quality control and DNA sequencing

One hundred nanograms of DNA was sheared to 550-700 bp using the Covaris LE220 and size selected using SPRI beads (Beckman Coulter). Ten nanograms of DNA for low input samples BottomLayer_2 and OS65 were sheared to 300 bp using the same selection. Libraries were created using the KAPA-Illumina library creation kit (KAPA Biosystems). Five cycles of PCR were performed for low input samples. The libraries were quantified using the KAPA Biosystems’ next-generation sequencing library qPCR kit and a Roche LightCycler 480 real-time PCR instrument. Sequencing was performed following a 2X151 indexed run recipe on the Illumina NovaSeq sequencer using NovaSeq XP V1 reagent kits and a S4 flowcell.

### Assembly, binning, taxonomic identification, and pan-genome analysis

As part of the JGI pipeline, for each sample, quality control of reads was performed with bbtools 38.26^78^ followed by assembly with spades 3.12.0 ^79^. Contigs with lengths less than 200 bp were removed. As part of the JGI pipeline, ORFs were called using the IMG pipeline (MAP v4.16.5). Briefly, this pipeline calls genes with Prodigal, GeneMark, and Infernal, and performs gene annotation using HMMER and LAST against Rfam 12.0, JGI’s custom RNA database, COG 2003, Pfam v30, and IMG-NR (exact version depending on date)^80–86^.

Following sample-specific assembly, reads were mapped to assembled contigs to create a cross-alignment using bowtie2 (v2.3.5.1)^87^. Binning was performed on the cross-aligned dataset using MetaBAT 2 ^29^ to create metagenome assembled genomes (MAGs). MAGs were assigned to the best hit in the GTDB-tk (r89) database^31,32^. MAGs belonging to the same phylum were used as input in the anvi’o ^30^ pangenome pipeline using default parameters. During the anvi’o pangenome pipeline, genes were called with Prodigal, annotated with COG and KO databases, and grouped into ortholog groups. Pan-genome taxa were manually selected from each phylum based on the similarity of ortholog groups. Taxa consisting of single MAGs or only poor-quality MAGs were removed from subsequent analyses. Additional analyses were performed on the anvi’o generated annotations unless otherwise noted.

The quality of the MAGs was assessed using the Genome Standards Consortium’s MIMAG (Minimum Information about a Metagenome-Assembled Genome) classification ^88^ (**Supplementary Data 2**).

### Comparison of pan-genome MAGs to prior MAGs

For the Chloroflexota MAGs that were identified to the family-level by GTDB-tk and the active Armatimonadota, we used fastANI (v0.1.3)^33^ on Kbase^89^ to compare a subset of MAGs from this study to the MAGs generated from the Mushroom Spring undermat^5,6^. These undermat MAGs were downloaded from RAST ^90^ on 2023/11/03 by logging in using a guest account and searching for Owner: Thiel, Vera. fastANI hits with greater than 94% identity were taken to suggest our MAGs were closely related to the Thiel MAGs and we used the more specific name given by Thiel^5,6^ (**Supplementary Data 2, 3)**.

### Metagenome statistical tests

The Shannon index was calculated using scikit-bio (v0.6)^91^ (**Supplementary Data 5)**. The Kruskal-Wallis test was implemented in scipy(v1.9.3). Bray-Curtis beta-diversity metric on relative metagenome abundance was calculated using scikit-bio (**Supplementary Data 6)**. ANOSIM (scikit-bio) was performed on the Bray-Curtis dissimilarity. nMDS of the Bray-Curtis dissimilarity was performed with scikit-learn (v1.3.2) with 2 dimensions (**Supplementary Fig. 2b**). ANCOM^40^ implemented in scikit-bio (v0.6) with ANOVA one-way significance testing was used to evaluate the differential abundance of genera between sample types (**Supplementary Data 7, Supplementary Fig. 3)**.

### Library preparation, quality control and sequencing (RNA)

rRNA was removed from 35.71 ng of total RNA using the Ribo-Zero(TM) rRNA Removal Kit (Epicentre). The rRNA depleted RNA was fragmented and reverse transcribed using random hexamers and SSII (Invitrogen) followed by second strand synthesis. Stranded cDNA libraries were generated using the Illumina Truseq Stranded RNA LT kit. Library-quality control and sequencing were performed using the same parameters as were used for generating the DNA libraries.

### Alignment, counting and normalization of RNA reads to metagenome assembled genomes

Bbtools 38.75 was used to perform trimming, filtering, and quality control^78^. Bowtie2 (v2.3.5.1) was used to align the quality-controlled RNA reads to 830 MAGs ^87^. Between 82-89% of reads that passed QC filtering aligned to the MAGs (**Supplementary Data 1**, doi:10.6084/m9.figshare.26530315). HTSeq-count ^92,93^ was used to count read pairs falling within predicted ORFs with flags (-s reverse -a 0). We set aqual = 0 in HTSeq-count due to the expected behavior of reads mapping to multiple locations in the total MAG set due to the presence of conserved regions across MAGs from the same pan-genome. 11-28% of reads were counted as properly aligning to a CDS (**Supplementary Data 1**, doi:10.6084/m9.figshare.26530315). The mapped counts to orthologous gene clusters identified for each pan-genome genus-level taxon were summed across all MAGs in the pan-genome (**Supplementary Data 8**). Gene clusters with low counts across all samples were removed prior to library normalization using edgeR (v3.30.3) ^94^ filterByExpr default settings with the exception min.count=1. Normalization was set with the “RLE” option^95^. A quasi-likelihood negative binomial generalized log-linear model (glmQLfit) fit to a natural spline (spline v 4.0.2) with three degrees of freedom was performed to estimate dispersion independently for each time series. Counts per million (CPM) were calculated from the normalized counts (**Supplementary Data 9**). For convenience of plotting, a single annotation was randomly selected from each gene cluster to use in the analyses (doi:10.6084/m9.figshare.26530315).

### Co-expression analysis by consensus Weighted Gene Co-expression Network Analysis (WGCNA)

A signed consensus WGCNA (v1.69) ^60,61^ was performed on gene clusters with mean CPM >1 that were log2(x+ 1) transformed, with each time series encoded as a separate experiment.

Individual co-expression networks were computed using the function blockwiseIndividualTOMs using a signed Pearson correlation. Soft-thresholding power (=18) was chosen as recommended in the WGCNA FAQ due to poor fit of scale-free topology. The output from this function was used as input in blockwiseConsensusModules to detect modules and compute eigengenes with the following changes from default: minModuleSize =30, deepSplit = 2, pamRespectsDendro = False, mergeCutHeight = 0.25 **(Supplementary Data 12**).

### Overrepresentation analysis

Gene clusters without a COG (Clusters of Orthologous Genes) Category^54^ were excluded in this analysis (32% of annotated gene clusters, and 8-22% of expressed gene clusters) (**Supplementary Data 10**). Overrepresentation analysis (ORA) was implemented in python 3.9 using a hypergeometric test (scipy.stats.hypergeom.cdf() (scipy v1.9.3)) and FDR multiple testing correction (statsmodels.api.stats.multipletest(method-=”fdr_bh”) v.0.13.5) for each genus. Only adjusted q-values < 0.05 are reported as significant (**Supplementary Data 16**).

### Armatimonadota phylogenetic tree

Annotated Armatimonadota genomes were downloaded from NCBI Genome on 2023/12/13. Sequences redundant with GTDB-tk (R207) were removed. Gtdb-tk de_novo_wf was used to create a concatenated single copy protein tree of the NCBI genomes, Thiel MAGs^5,6^, and MAGs from this study, and then decorated with Armatimonadota and Chloroflexota genomes from the GTDB-tk database^31,32^. Tree visualization was performed in iTOL^96^ (**Supplementary Fig. 10**).

### Armatimonadota genomic potential

A variety of approaches were used to interpret the metabolic potential of the selected Armatimonadota MAGs. KEGG orthology numbers (KO) were extracted from the JGI/IMG annotations and combined into a concatenated KO list per genus for upload to KEGG Reconstruct Pathway^97^ (accessed 2022/10/12, 2022/10/22). We also examined the annotations created in the anvi’o pipeline, and another alternative annotation using RASTtk on KBase ^89,90,98,99^. Some sequences were used as input to BLASTP to annotate NCBI conserved domains as additional interpretation^100,101^.

JGI predicted ORFs for the Armatimonadota MAGs were imputed into specific pipelines to predict different metabolic properties. The GapMind^102^ webserver (accessed 2022/09/13-2022/09/28) was used for carbon source prediction. The summary output was captured for each MAG and manually concatenated to determine the predicted phenotype for each genus (**Supplementary Data 18)**. The dbCAN2 web server ^103,104^ (accessed 2022/10/11-2022/10/18) was used for carbohydrate active enzyme (CAZyme) prediction using the options: HMMER (dbCAN), DIAMOND (CAZy), and HMMER (dbCAN-sub) search. Genes detected as CAZymes with 2 or more methods were examined further for their function and reported as the total number of genes per CAZy category per genome and normalized by the total number of genes in each MAG (**Supplementary Data 19)**. For iron metabolism and extracellular electron transfer, FeGenie (v1.0) (options --orfs --all_results --heme) ^105^ was run (**Supplementary Data 20, 21**). Genes containing heme-binding domains were run through the NCBI conserved domain database^101^ and interproscan^106,107^ webservers to look for domains of interest (accessed September 2023). Putative operon context on assembled contigs was also considered in interpreting function.

### Hydrogenase classification

To classify the annotated hydrogenases, sequences annotated as hydrogenase were uploaded to HydDB (accessed 2023/03/09)^108^. Only catalytic subunits are classified by the webserver, and other subunits entered resulted in a “NONHYDROGENASE” result (**Supplementary Data 22**).

## Supplementary Information

**Description of Additional Supplementary Files**

**Supplementary Data 1-22**

**Reporting Summary**

## Data availability

Raw sequence reads can be found on the SRA at SRP191317, SRP191318, SRP191319, SRP191321, SRP191322, SRP191324, SRP191332, SRP191328, SRP191329, SRP191333, SRP213393, SRP213396, SRP213395, SRP213397, SRP213394, SRP191337, SRP213390, SRP213392, SRP213391, SRP191334, SRP213388, SRP191350, SRP213398, SRP213402, SRP239941, SRP213411, SRP213409, SRP213410, SRP213412, SRP213413, SRP213403, SRP213408, SRP213407, SRP213406, SRP259901, SRP259948, SRP259967, SRP260043, SRP259966, SRP260069, SRP260079, SRP260124, SRP260130, SRP260133, SRP260135, SRP260129, SRP288870, SRP288866, SRP288863, SRP288864, SRP288862, SRP288861, SRP288868, SRP288867, SRP288819, SRP288812, SRP288826, SRP288804, SRP288815, SRP288817, SRP288820, SRP288824, SRP288827, SRP288823, SRP288853, SRP299000.

NCBI Bioprojects associated with the samples are: PRJNA518216, PRJNA518217, PRJNA518218, PRJNA518219, PRJNA518220, PRJNA518221, PRJNA518222, PRJNA518223, PRJNA518224, PRJNA518225, PRJNA539609, PRJNA539610, PRJNA539611, PRJNA539612, PRJNA539608, PRJNA518227, PRJNA539605, PRJNA539606, PRJNA539607, PRJNA518226, PRJNA539604, PRJNA518228, PRJNA539613, PRJNA539614, PRJNA571079, PRJNA539619, PRJNA539620, PRJNA539621, PRJNA539622, PRJNA539623, PRJNA539615, PRJNA539618, PRJNA539617, PRJNA539616, PRJNA621571, PRJNA621572, PRJNA622232, PRJNA621573, PRJNA621574, PRJNA621575, PRJNA621576, PRJNA621577, PRJNA621578, PRJNA621580, PRJNA621581, PRJNA621579, PRJNA653764, PRJNA653765, PRJNA653766, PRJNA653767, PRJNA653768, PRJNA653769, PRJNA653770, PRJNA653771, PRJNA653752, PRJNA653753, PRJNA653754, PRJNA653755, PRJNA653756, PRJNA653757, PRJNA653758, PRJNA653759, PRJNA653760, PRJNA653761, PRJNA653762, PRJNA677357.

JGI/IMG portal accessions can be found in **Supplementary Data 1.**

MAG sequences and annotations, and some additional data used in plotting have been deposited at Figshare: doi:10.6084/m9.figshare.26530315

Analyzed datasets are provided as Supplementary Data.

## Code availability

Custom scripts used to generate and analyze data can be found in our repository on GitHub: https://github.com/carnegie/YNP_MSOS_metagenomics_metatranscriptomics

## Author contributions

D.B., F.B.Y., and A.N.S. conceived and designed the research. F.B.Y. and A.N.S. carried out the analyses. A.N.S. and D.B. wrote the initial manuscript. A.N.S., F.B.Y, D.B., and A.R.G. edited the manuscript.

## Supporting information

Description of Supplementary Data Files

## Acknowledgements

Sampling was performed under NPS Park Permits: YELL-5494 (to David Ward, multi year), YELL-5660 (to Devaki Bhaya, 2007-2008), YELL 5694 (to Devaki Bhaya 2007-2009). We thank Mary Bateson and all members of the Ward lab as well as Michael Kuhl, Sheila Jensen, Anne Steunou, Oliver Kilian, Melissa Adams and Michelle Davidson for their assistance in sampling over the years. We thank Rick Kim and Jackie Meisel for assistance with DNA and RNA extraction and metagenome binning protocols. We thank Vera Thiel, Zeqian Li, Seppe Kuehn, Chris Howe, Alison Smith, Stephen Rowden, Andre Holzer, and members of the Bhaya, Grossman, Burlacot and Yeh labs for discussions. Library prep and sequencing was performed by JGI under proposal 503441. The work (proposal: 10.46936/10.25585/60001132) conducted by the U.S. Department of Energy Joint Genome Institute (https://ror.org/04xm1d337), a DOE Office of Science User Facility, is supported by the Office of Science of the U.S. Department of Energy operated under Contract No. DE-AC02-05CH11231. This work was supported by BBSRC-NSF/BIO #1921429 awarded to D. Bhaya and A. Grossman (Carnegie Science) and NSF#2125965 (Emerging Frontiers) to Devaki Bhaya (Carnegie Science).

