## Supplementary Figures 1-10 for "Abundant and active community members respond to the diel cycle in hot spring phototrophic mats"

**This file contains:**
Supplementary Figures 1-10
**Other Supplementary Material includes the following:**
Supplementary Data 1-22

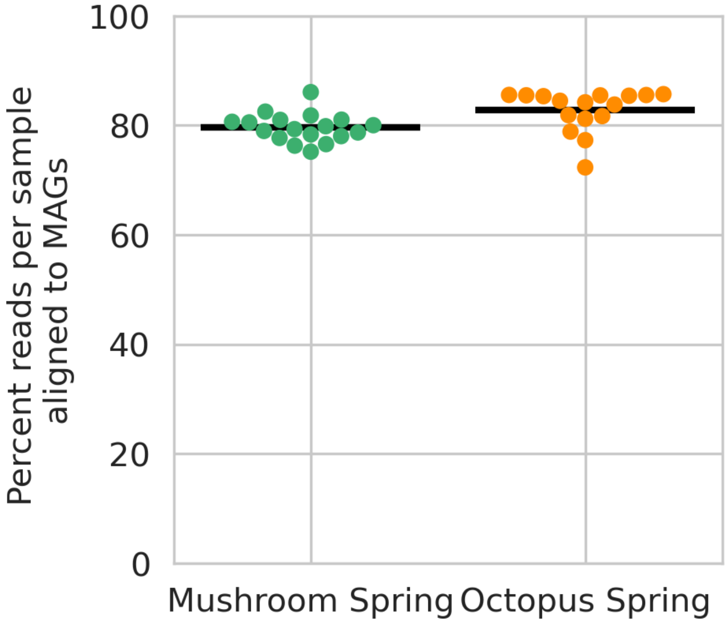


**Supplementary Figure 1: Percent of reads mapping to MAGs**
Alignment of metagenome paired reads to the 830 selected MAGs for all 34 metagenome samples. Mean percent reads aligned per spring are indicated with the black line.

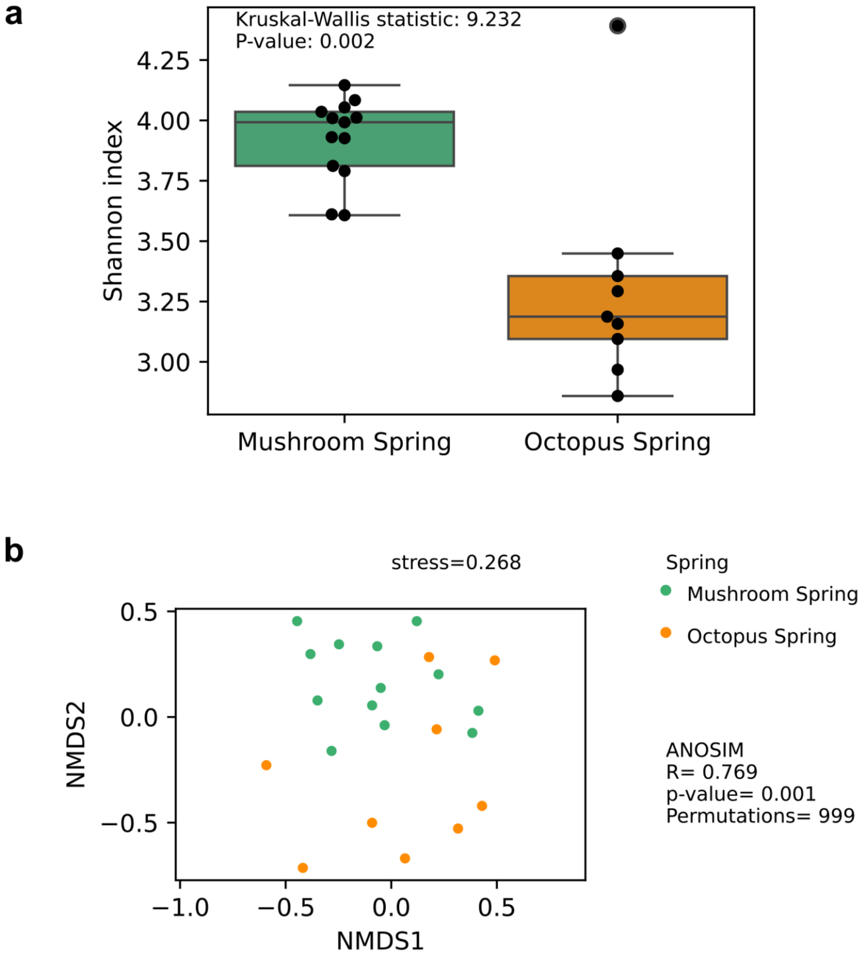


**Supplementary Figure 2: Diversity metrics of 60°C samples**
**(a). Shannon alpha diversity index on metagenome relative abundance.** Boxplots show the median, upper and lower quartiles of the Shannon index. Whiskers are 1.5 times the interquartile range, and points are those falling outside the whiskers. **(b). Bray-Curtis dissimilarity of the metagenome relative abundance.** 2-D nMDS on Bray-Curtis dissimilarity is plotted. The ANOSIM test was performed to compare Mushroom Spring and Octopus Spring, total samples =22.

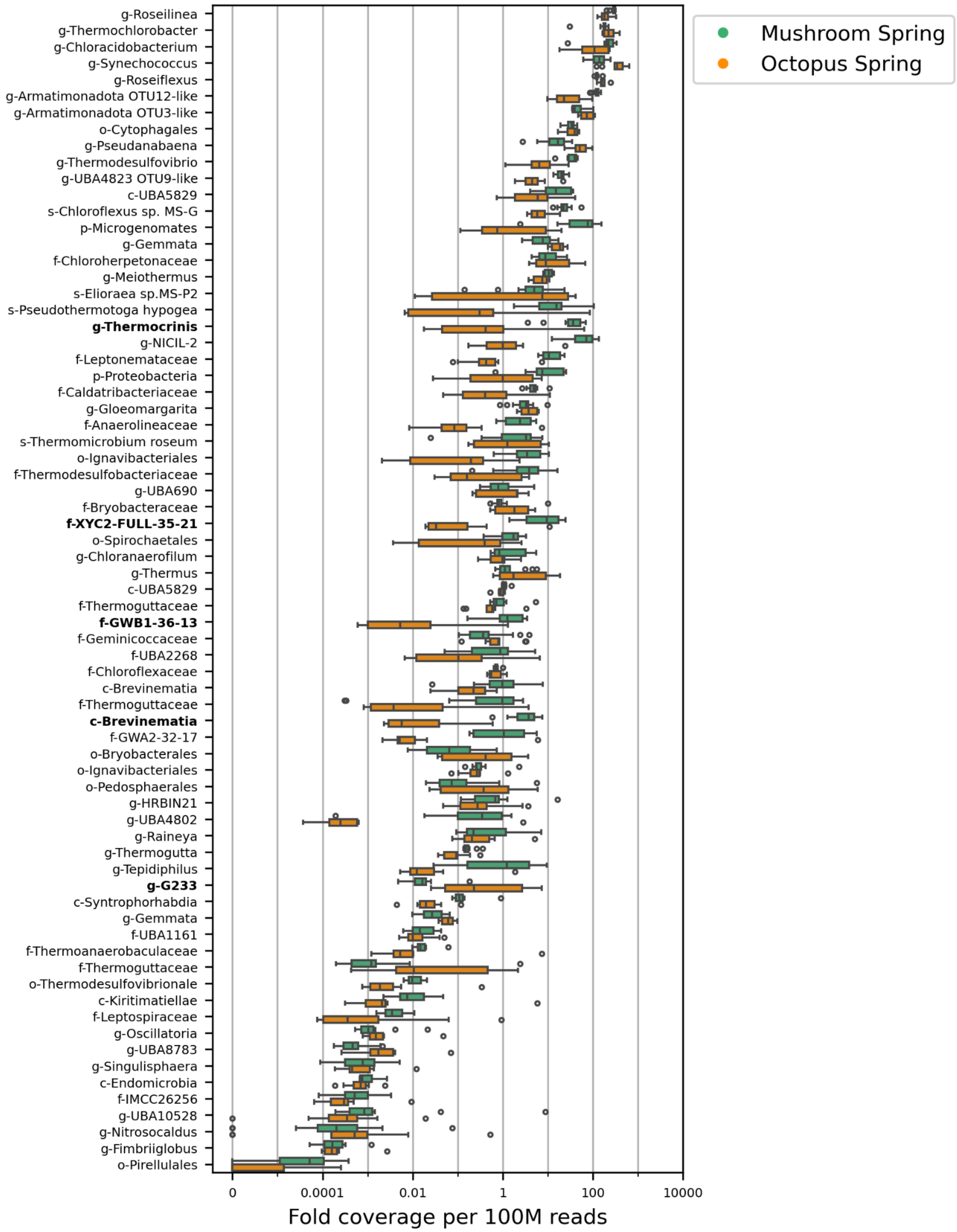


**Supplementary Figure 3: Summary of metagenome relative abundance at 60°C**
The median, upper and lower quartile of the fold coverage of each pan-genome lineage are depicted in the box plot. Whiskers are 1.5 times the interquartile range, and points are those falling outside the whiskers. **Bold** = significantly different relative abundance by the ANCOM test between springs at 60°C.

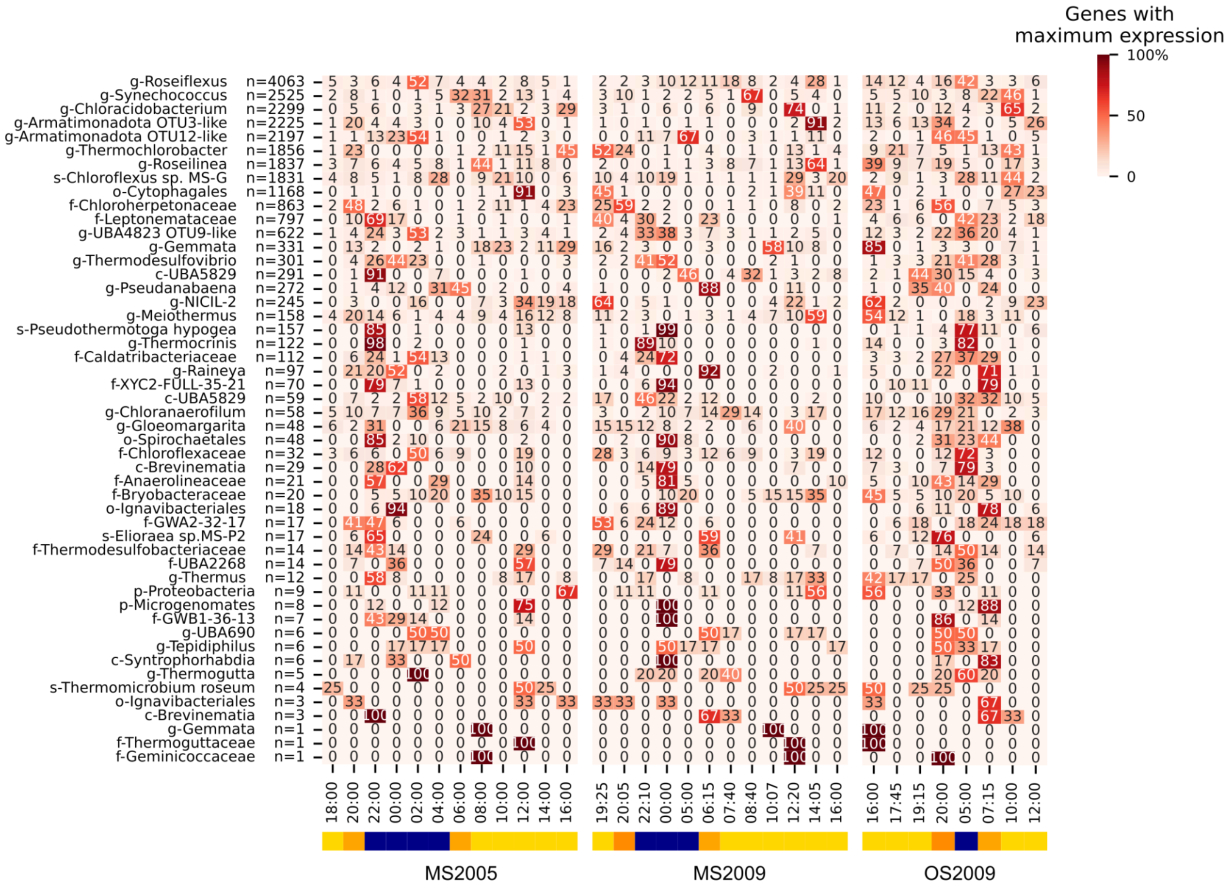


**Supplementary Figure 4: Percent of highly expressed genes per taxon with maximum expression at each timepoint for all three metatranscriptome time series.**

Heatmap of percent of highly expressed gene clusters (mean CPM >1) per taxon that had maximum expression at each time point. n= number of genes analyzed per taxon. Only taxa with at least one highly expressed gene are shown. Bars at the bottom indicate qualitative time of day: yellow: day, orange: dusk or dawn, blue: night (Note that sampling in MS2005 and MS2009 has more time points than OS2009).

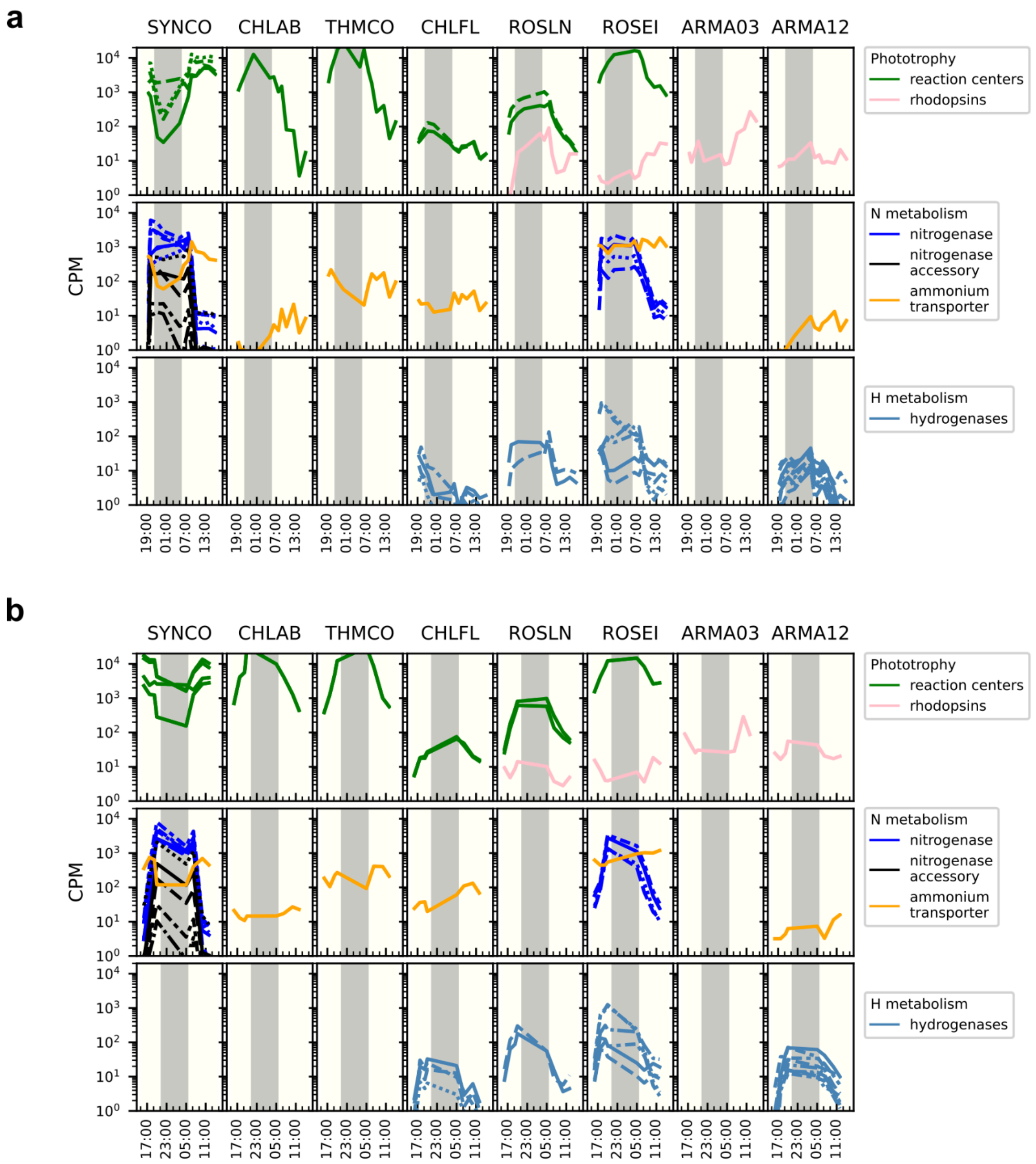


**Supplementary Figure 5: Response of sentinel pathways to the diel cycle in the eight most active taxa in MS2009 and OS2009 datasets**

Gene expression of the same sentinel genes from **Fig. 4** are plotted from the (**a)** MS2009 (**b)** OS2009 metatranscriptome for the eight active taxa to show similarities and differences between the time series. Panels without curves indicate that a gene containing that annotation was not expressed in that taxon. Gray bar: Night period. CPM: counts per million. Underlying data is found in **Supplementary Data 11.**

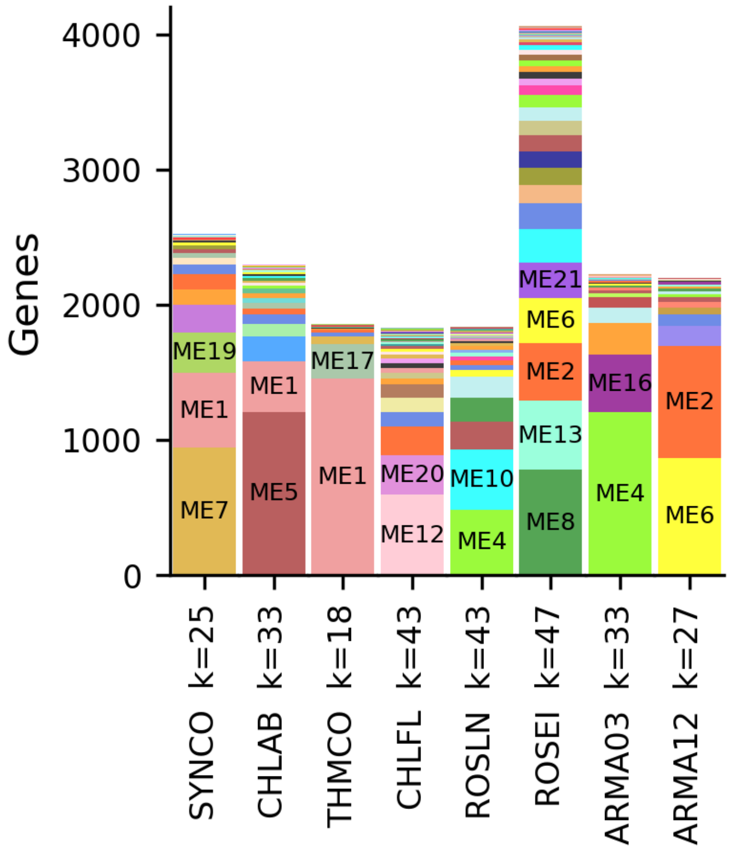


**Supplementary Figure 6: Distribution of genes in WGCNA modules in the eight most active taxa**

Alternative view of data in **Fig. 5a,** the number of highly expressed genes per taxon in each module. Modules with greater than 250 genes per genus are labeled. k=number of WGCNA modules per taxon.

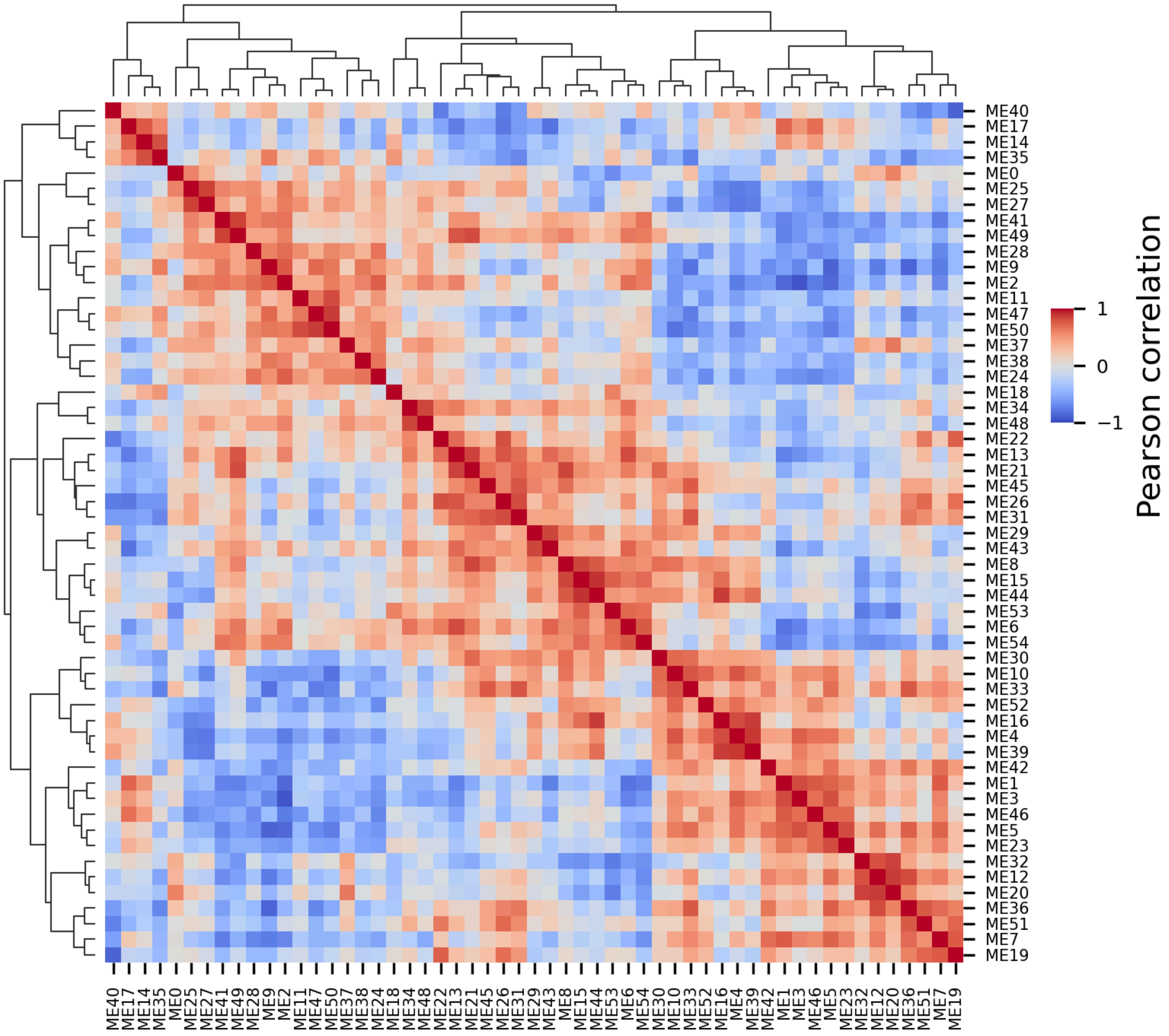


**Supplementary Figure 7: Pearson correlation of WGCNA module eigengenes**
The Pearson correlation and hierarchical clustering was used for ordering the WGCNA modules in **Fig. 5b** and **Fig. 6**. Underlying data is found in **Supplementary Data 14.**

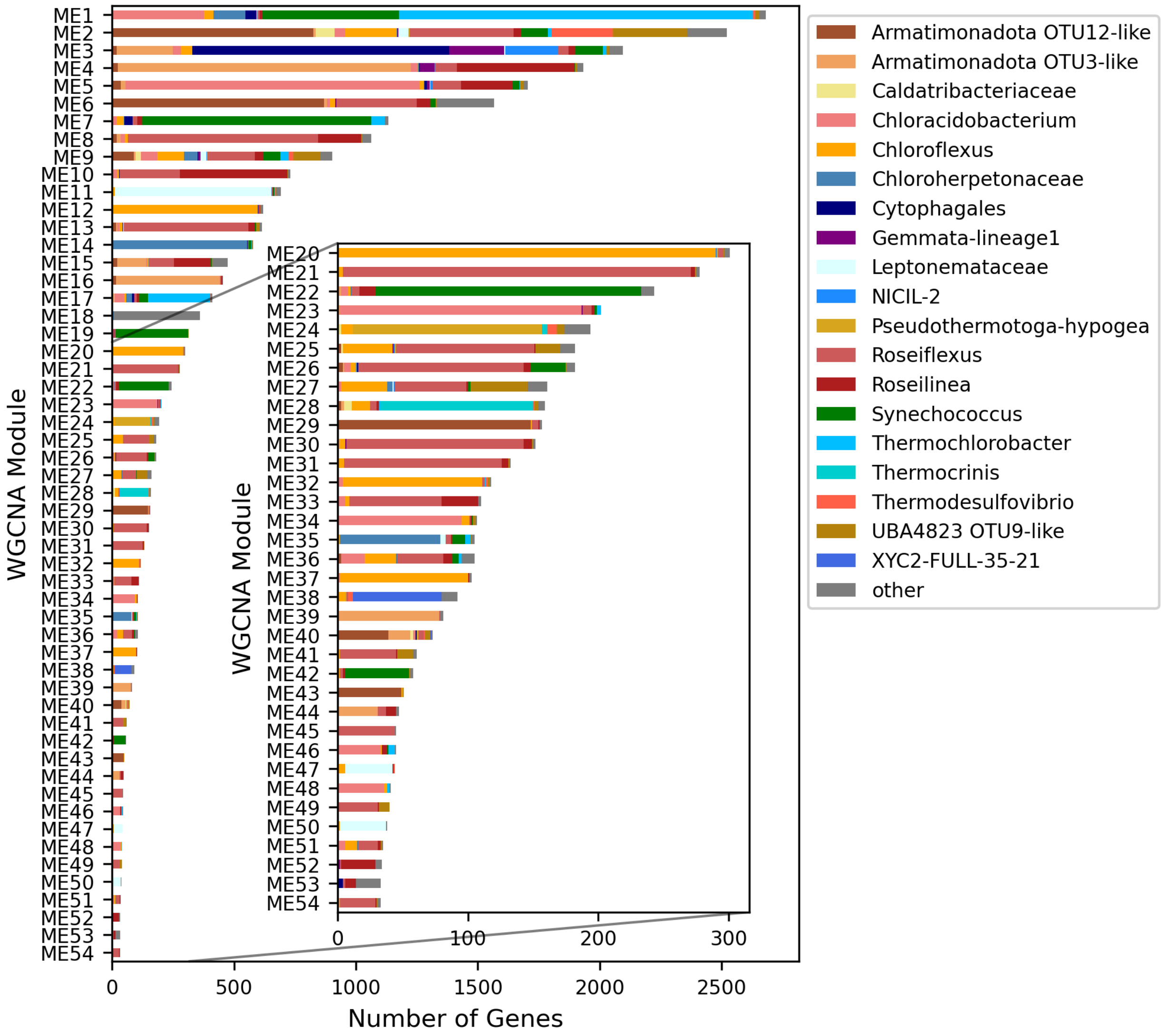


**Supplementary Figure 8: Taxa composition of WGCNA modules**

Taxa composition of genes in consensus WGCNA modules identified in highly expressed genes in all three time series (**Fig. 5**). An inset shows more detail for smaller WGCNA modules.

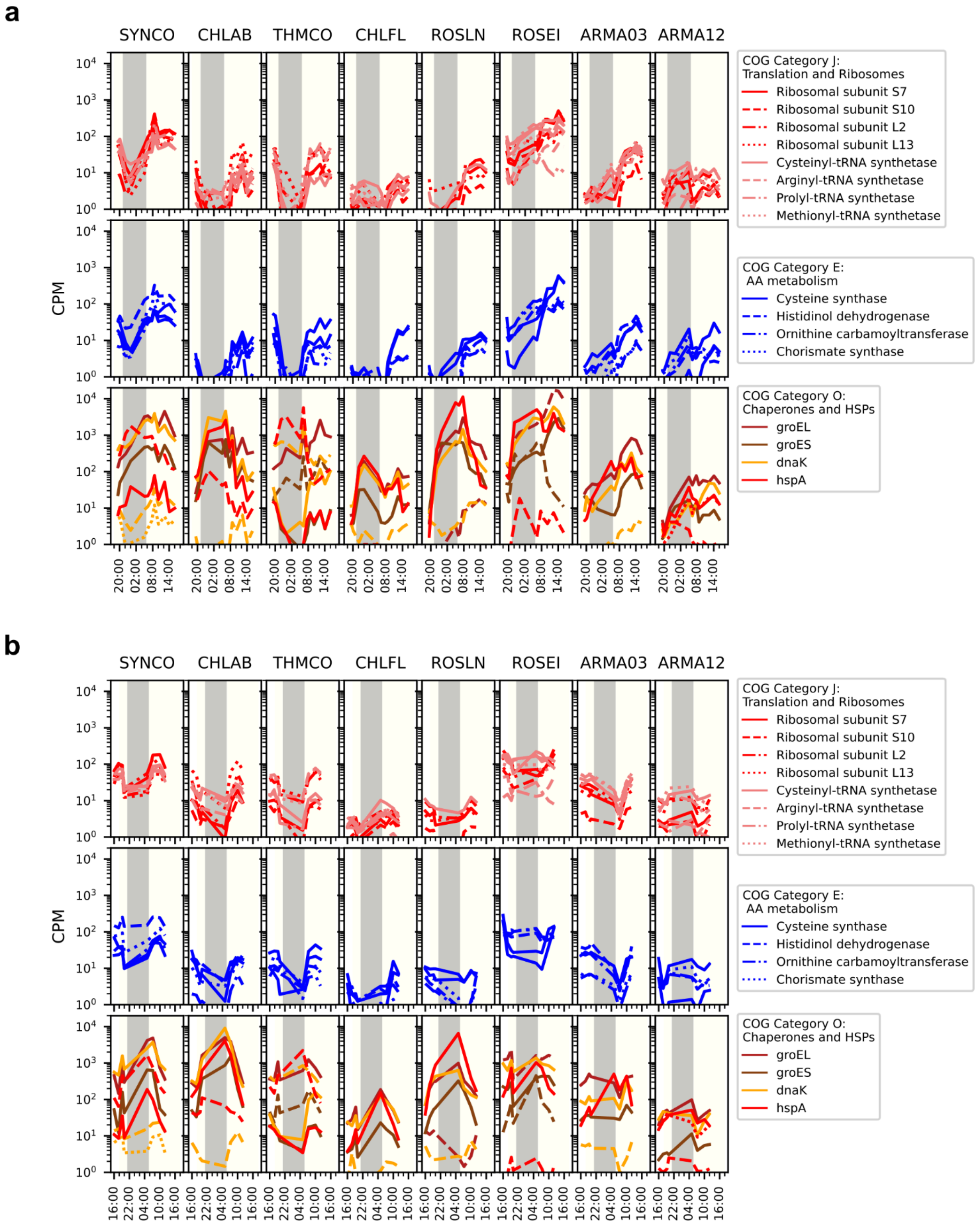


**Supplementary Figure 9: Expression patterns of selected genes from the COG Category overrepresentation analysis in MS2009 and OS2009 datasets**

Gene expression of the same sentinel genes from **Fig. 7** are plotted from the (**a**)MS2009 and (**b**) OS2009 metatranscriptome for the eight active taxa to show similarities and differences between the time series. Each line is a different gene. Top: Category J, Translation and Ribosomes. Middle: Category E, Amino acid (AA) metabolism. Bottom: Category O, Chaperones and heat shock proteins (HSPs). CPM: counts per million. Underlying data is found in **Supplementary Data 17**.

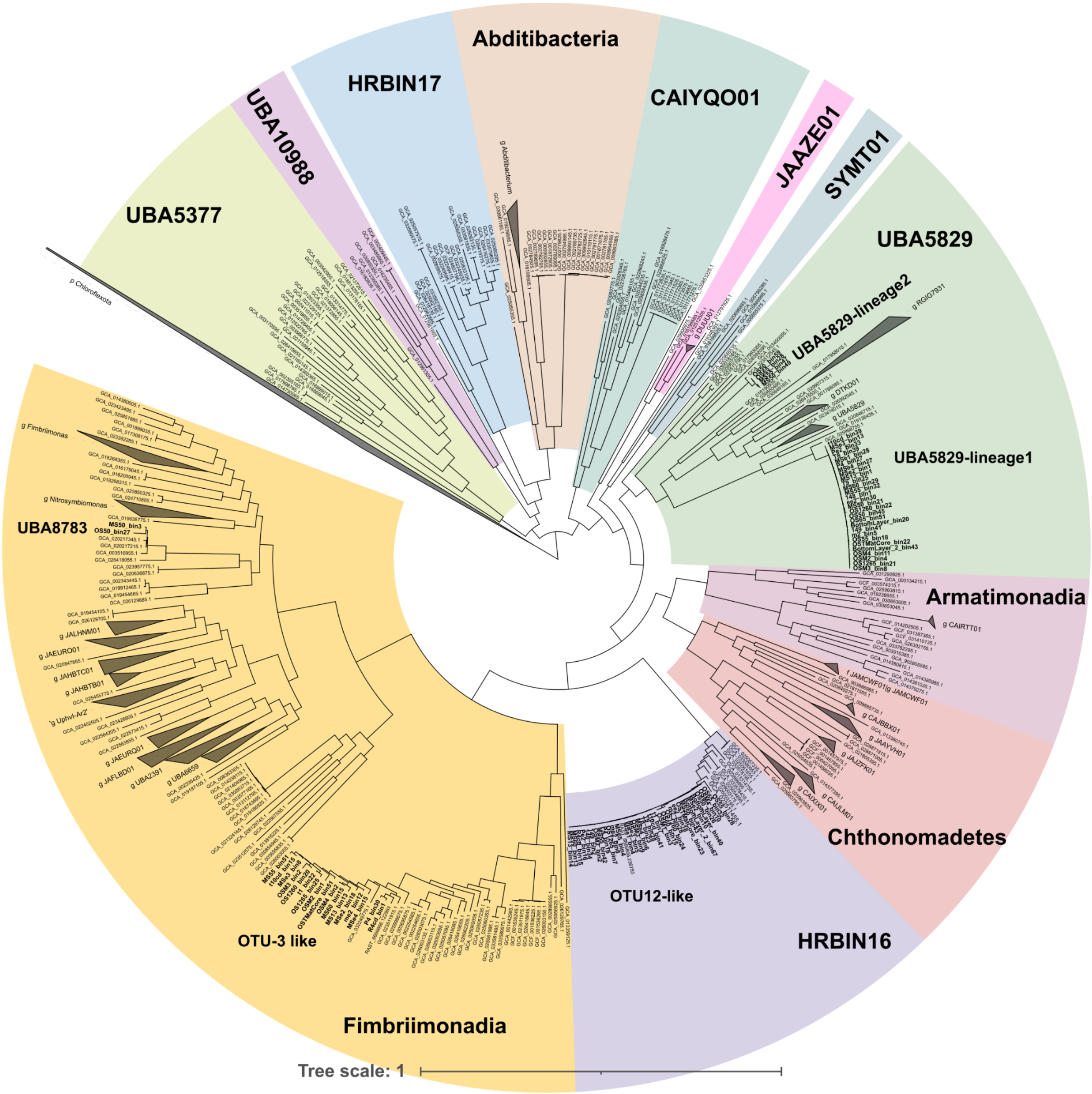


**Supplementary Figure 10: Armatimonadota MAGs in context with available sequences from GTDB and NCBI**
A concatenated single copy protein tree containing GTDB (R207) reference genomes, genomes classified as Armatimonadota on NCBI (downloaded 2023/12/13), the two MAGs from Thiel 2016, 2017, and the Armatimonadota MAGs from this study (bolded text) was created using GTDB-tk. Class assignments by GTDB-tk are indicated by colored backgrounds. Scale bar is amino acid substitutions per site.
