## Supplementary material for "Abundant and active community members respond to the diel cycle in hot spring phototrophic mats": Description of Supplementary Data Files

### Description of Additional Supplementary Files

**Supplementary Data 1: Sample accessions and metadata**

**Supplementary Data 2: MAG metadata**
Samples metadata draws from **Supplementary Data 1**. MAG metadata includes genome size, number of contigs, GC content, estimates of completeness and contamination from checkM, the fold coverage of the MAG in the sample it is from, number of annotations from JGI/IMG and anvi'o pipelines, whether the MAG was selected for further analysis, the pan-genome lineage designation and the full GTDBtk results per MAG. MIMAG classification is also included.

**Supplementary Data 3: fastANI results comparing selected MAGs**

**Supplementary Data 4: Relative abundance of pan-genome taxa in metagenome samples**

**Supplementary Data 5: Metagenome Shannon indices**

**Supplementary Data 6: Bray-Curtis dissimilarity matrix for metagenome samples**

**Supplementary Data 7: ANCOM results for spring-specific taxa in 60°C metagenomes**
The percentile columns show the relative metagenome coverage for that taxon in each spring at the different percentile values for the taxon. The W-statistic is the number of taxa that the taxon has been tested to be significantly different against.

**Supplementary Data 8: RNA mapping output from HTSeq-count aggregated per ortholog group in each pan-genome taxon group**

**Supplementary Data 9: Normalized RNA counts per million per times series using the pan-genome taxon ortholog groups**
Each sheet is a different sampling series

**Supplementary Data 10: Summary statistics on metatranscriptome counts per gene and annotation type**
The total ORFs includes hypothetical genes, "annotated" is only genes with a KOfam or COG annotation. For "detected" columns, this represents at least one read per gene, and "highexpress" represents a mean CPM > 1 in that time series.

**Supplementary Data 11: Selected expression data of sentinel genes for Figure 4 and associated Supplementary Figure 5.**
The annotation is given for both COG and KOfam accessions. Sample names correspond to those in **Supplementary Data 1.**

**Supplementary Data 12: WGCNA module assignment to genes**

**Supplementary Data 13: Eigengene expression values**

**Supplementary Data 14: Pearson Correlation of WGCNA module eigengene expression patterns**

**Supplementary Data 15: Range of expression values (CPM) for each pan-genome taxon observed in all 32 metatranscriptome samples.**

**Supplementary Data 16: Overrepresentation analysis results for COG Categories in WGCNA modules in pan-genome taxa.**

**Supplementary Data 17: Selected expression data for genes for Figure 7 and Supplementary Figure 9.**
The annotation is given for both COG and KOfam accessions. Sample names correspond to those in **Supplementary Data 1.**

**Supplementary Data 18: GapMind carbon source prediction results on Armatimonadota OTU3-like and Armatimonadota OTU12-like.**

**Supplementary Data 19: dbCAN carbohydrate active enzymes prediction results on Armatimonadota OTU3-like and OTU12-like**

**Supplementary Data 20: FeGenie heme binding protein prediction results on Armatimonadota OTU3-like and Armatimonadota OTU12-like.**

**Supplementary Data 21: FeGenie results for other categories on Armatimonadota OTU3-like and Armatimonadota OTU12-like.**

**Supplementary Data 22: HydDB classification of genes annotated as “hydrogenase” in the eight most active taxa.**
Only the catalytic subunit is classified by the database into the different categories, other subunits give a “NONHYDROGENASE” result.
